# Hematopoietic lineages diverge within the stem cell compartment

**DOI:** 10.1101/2020.08.21.261552

**Authors:** Mina N. F. Morcos, Congxin Li, Clara M. Munz, Alessandro Greco, Nicole Dressel, Susanne Reinhardt, Andreas Dahl, Nils B. Becker, Axel Roers, Thomas Höfer, Alexander Gerbaulet

## Abstract

Hematopoietic stem cells (HSCs) produce a highly diverse array of cell lineages. To assay hematopoietic differentiation with minimal experimental perturbation, non-invasive methods for heritable labeling^1–3^ or barcoding^4–7^ of HSCs *in vivo* have recently been developed and used to study lineage fate of HSCs in physiological conditions. However, the differentiation pathways leading from HSCs to mature cells remain controversial^8^, with suggested models ranging from gradual lineage restriction in a branching cascade of progenitors to HSCs already making ultimate lineage decisions. Here we show, by iterating HSC fate-mapping, mitotic history tracking, single-cell RNA-sequencing and computational inference, that the major differentiation routes to megakaryocytes, erythro-myeloid cells and lymphocytes split within HSCs. We identify the hitherto elusive self-renewing source of physiological hematopoiesis as an HSC subpopulation co-expressing high levels of Sca-1 and CD201. Downstream, HSCs reduce Sca-1 expression and enter into either thrombopoiesis or erythro-myelopoiesis, or retain high Sca-1 levels and the ability to generate lymphocytes. Moreover, we show that a distinct population of CD48^−/lo^ megakaryocyte progenitors links HSCs to megakaryocytes. This direct thrombopoiesis pathway is independent of the classical pathway of megakaryocyte differentiation via multipotent progenitors and becomes the dominant platelet production line upon enhanced thrombopoietin signaling. Our results define a hierarchy of self-renewal and lineage decisions within HSCs in native hematopoiesis. Methodologically, we provide a blueprint for mapping physiological differentiation pathways of stem cells and probing their regulation.

---

Immuno-phenotypic HSCs (lineage^−^ Sca-1^+^ CD117^+^ (LSK) CD48^−/lo^ CD150^+^; Extended Data Fig. 1) are a heterogeneous population with regard to proliferative activity, lineage fate and repopulation capacity after transplantation; in particular, numerous HSC transplantation studies have described, to varying degrees, lineage fate biases of HSCs (reviewed in ^9^). The production rate of mature cells from HSCs is massively enhanced after transplantation compared to physiological conditions^4^, and it remains unknown whether lineage differentiation pathways also differ between the two settings. Recent *in vivo* barcoding studies of HSCs in native hematopoiesis have revealed HSC clones with different lineage output^5,6,10^, which appear to resemble those seen after HSC transplantation^11,12^. These include HSCs producing only megakaryocyte progenitors (MkPs) independent of multipotent progenitors (MPPs)^10,11,12^. However, how distinct HSC and progenitor subpopulations are related to each other and at what level they commit to lineage pathways in native hematopoiesis is unknown (Fig. 1a). Specifically, the observation of HSCs feeding exclusively into thrombopoiesis, which appear to be distinct from the HSCs supplying the thrombopoiesis pathway via MPPs^13,14^ and committed myeloid progenitors^15^, raise the question of which lineage pathway produces platelets during native hematopoiesis. These questions are intimately linked with another controversy that has arisen in the context of the recent HSC fate mapping studies, suggesting steady but infrequent input from HSCs to hematopoiesis^2,4^ or more active contribution^3,16,17^. We reasoned that integrating independent data on HSC differentiation, proliferation and molecular heterogeneity, all obtained under physiological conditions, will help resolve these fundamental open questions.

**Figure 1.**
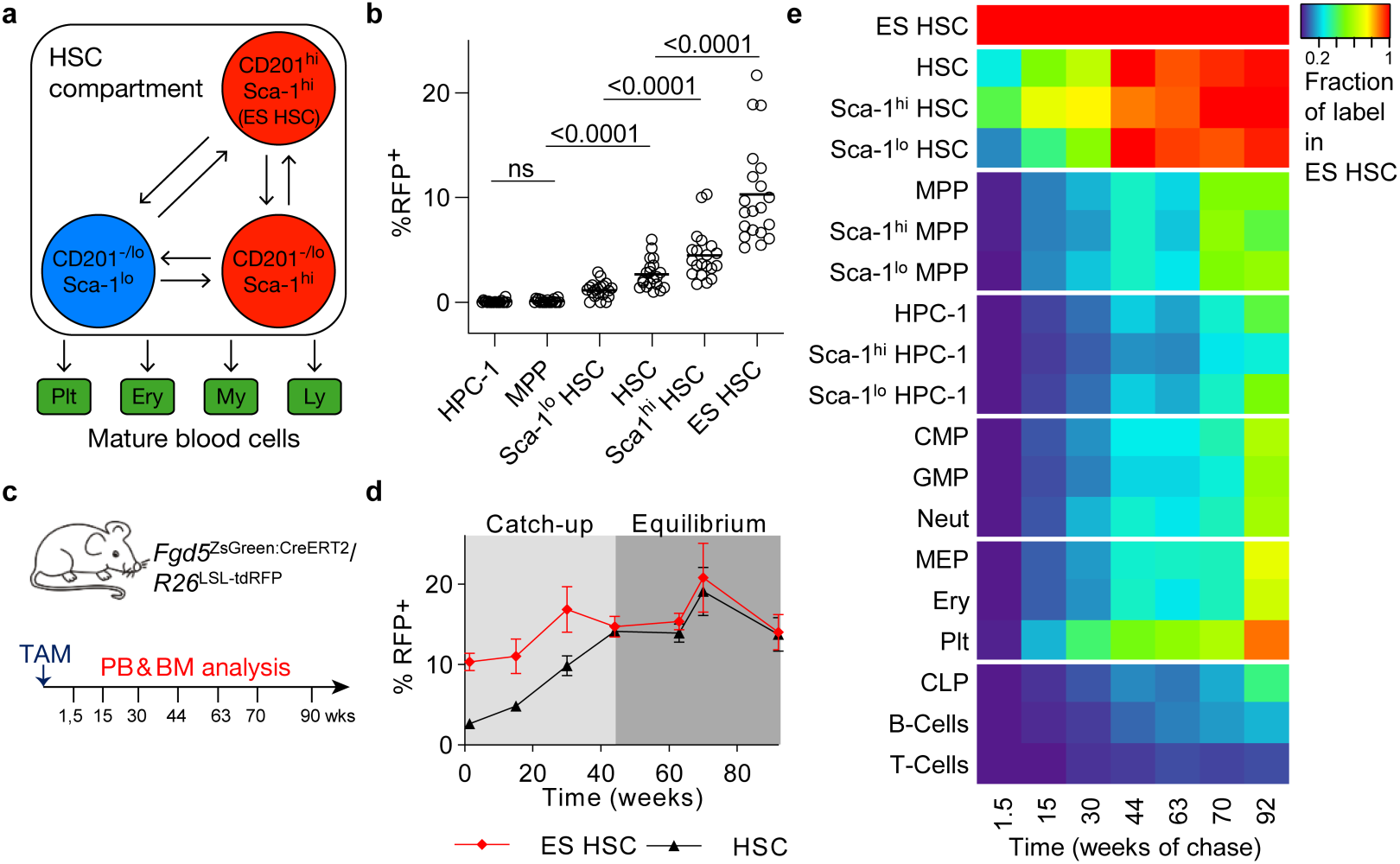
Fate mapping of CD201^hi^ Sca-1^hi^ HSCs. **a**, Subdivision of the immuno-phenotypic HSC population (LSK CD48^-/lo^CD150^+^) by CD201 (EPCR) and Sca-1 expression; ES-HSC, CD201(EPCR)^hi^ Sca-1^hi^ HSCs. Relations between HSC populations in native hematopoiesis and how they connect to lineage pathways will be studied. **b**, Bone marrow populations of *Fgd5*^ZsGreen:CreERT2^*/R26*^LSL-tdRFP^ mice were analysed 10 - 11 days after TAM induction for RFP expression (n=20, 4 independent experiments, one-way ANOVA with Sidak correction); for flow cytometry phenotypes see Extended Data Fig. 1a. **c-e**, To map HSC fate mapping over time, *Fgd5*^ZsGreen:CreERT2^*/R26*^LSL-tdRFP^ mice (n=82) were TAM-induced, groups of mice were sacrificed at indicated time points, and bone marrow populations and peripheral blood leukocytes were analysed for RFP expression (pooled data from 5 independent experiments are shown). **d**, Percentage of RFP^+^ HSCs and ES HSCs (mean and SEM). **e**, Fractions of labeled peripheral blood and bone marrow cells relative to ES HSCs at indicated time points; the labeling frequencies of the cell populations from each individual were normalized to its ES HSC labeling at final bone marrow analysis.

## CD201^hi^ Sca-1^hi^ HSCs are the source of physiological hematopoiesis

In previous HSC fate mapping studies, the actual stem cell source that supplies hematopoiesis over the lifetime of the mouse, called tip HSCs, has remained elusive^2,3,16,17^. The defining criterion for such tip HSCs is that after label induction, constant labeling frequency is maintained by perfect self-renewal^18^. In existing HSC fate mapping data, however, a several-fold increase of HSC labeling frequency over at least one year has been observed that is fueled by an unknown subpopulation of tip stem cells within the phenotypic HSCs^2,3,16,17^. To selectively label HSCs for fate mapping, we generated the *Fgd5*^ZsGreen:CreERT2^*/R26*^LSL-tdRFP^ mouse model (Extended Data Fig. 2a), which combines an inducible Cre expressed in HSCs^1^ (Extended Data Fig. 2b), with an excision reporter with high activation threshold^19,20^. Indeed, tamoxifen (TAM) application labeled exclusively HSCs (Fig. 1b, Extended Data Fig. 2c-e), thus minimizing the problem of concomitant labeling of progenitors downstream of HSCs seen in previous studies^3,16,17^. To characterize HSC heterogeneity, we used surface levels of CD201 (also termed EPCR) and Sca-1, as either marker has previously been shown to correlate with repopulation potential of transplanted HSCs^21–25^. RFP-labeled cells were strongly enriched among CD201^hi^ Sca-1^hi^ HSCs (ES HSC) (Fig. 1b). Indeed, ten sorted RFP^+^ HSCs, but not RFP^−^ HSCs, isolated from recently labeled *Fgd5*^ZsGreen:CreERT2^*/R26*^LSL-tdRFP^ mice, yielded robust multi-lineage repopulation in a competitive transplantation assay (Extended Data Fig. 2f-h).

We induced a large cohort (n=82) of *Fgd5*^ZsGreen:CreERT2^/*R26*^LSL-tdRFP^ mice to follow label propagation over time (Fig. 1c). Similar to previous studies^2,3,16,17^, labeling frequency of the total HSC population increased several-fold over the first year (Fig. 1d, black line). By contrast, labeling of ES HSCs remained approximately constant (Fig. 1d, red line, Extended Data Fig. 2i, j), which indicates perfect self-renewal of this HSC subpopulation. Moreover, the labeling frequency of all HSCs equilibrated to ES HSCs, implying that the latter are the ultimate source of all HSCs. Taken together, these data identify ES HSCs as the tip stem cells of physiological hematopoiesis.

To quantify label propagation from ES HSCs through the hematopoietic system, we calculated the labeling frequencies of downstream bone marrow and peripheral blood populations relative to ES HSC labeling at final analysis of a given individual, thus normalizing for variability of initial RFP labeling frequency in ES HSCs (Extended Data Fig. 2i, j). All progenitor and mature cell populations slowly increased their labeling frequencies, trailing HSCs. Labeling of MPPs increased with considerable delay and did not equilibrate with HSC labeling within 21 months of chase (Fig. 1e), which implies that progression of cells from ES HSCs to MPPs represents a kinetic bottleneck. In mature cells, labeling frequency increased fastest in platelets, slowest in T cells and at intermediate rate in granulocytes and erythrocytes. Thus, label induced in ES HSCs reached all major progenitors and mature cells of the hematopoietic system, but with different kinetics.

## Combining ES HSC fate mapping with mitotic tracking yields lineage pathways

We asked whether the detailed and time-resolved HSC fate mapping (Fig. 1e) data contain information on lineage pathways. Clearly, label from ES HSCs arrived in progenitors before it entered mature cells (e.g., common-lymphoid progenitors (CLPs) and B cells, respectively, in Fig. 1e). Applying this principle to our data ruled out certain pathways (specifically, classically defined MEPs could not be precursors of platelets, as platelets were more strongly labeled than MEPs from 15 weeks of chase onward; Fig. 1e). We now show that the data can be used to chart lineage pathways. The topology of lineage pathways emerging from stem cells is primarily defined by branch points; it may also contain convergence points when a downstream population has more than one progenitor (see below). Within a given pathway topology, three measurable quantities are closely related: the rates of *cell differentiation* and *proliferation* at each node define the *size of the cell population* constituting this node (Extended Data Fig. 3a). The *in vivo* values of differentiation rates, proliferation rates and population sizes can be measured by, respectively, HSC fate mapping^26^, dilution of fluorescently labeled histones (e.g., H2B-GFP^27^), and cell counting. We reasoned that obtaining these data for all cell populations in a stem cell differentiation pathway will also yield the pathway topology. To examine this, we generated a toy model for a generic branching lineage topology *in silico* (Extended Data Fig. 3b) and simulated fate mapping and mitotic tracking data. We then constructed all principally possible topologies linking stem cells, progenitors and mature cells (Extended Data Fig. 3c), and asked which topology is compatible with the simulated data. Statistical model selection based on the simulated data with added noise indeed recovered the true pathway and rejected all others (Extended Data Fig. 3d, e). By contrast, using the *in silico* fate mapping data alone did not single out the correct model (Extended Data Fig. 3f). Thus, lineage pathways downstream of ES HSCs can be inferred from combined data on stem cell fate mapping, mitotic history and cell numbers.

## The myeloid differentiation pathway diverges within HSCs

We applied this inference approach to our experimental fate mapping data, complementing them with measurements of H2B-GFP label dilution, and stem and progenitor cell numbers measured relative to lineage-negative (lin^−^) cells. In addition to subdivision of HSCs by CD201 and Sca-1 expression, we stratified MPPs and HPCs-1 by Sca-1 expression. This yielded a graph of all possible pathway topologies, with unidirectional differentiation steps (as implied by transplantation studies) and the possibility of reversible Sca-1 loss (Fig. 2a). We systematically enumerated 144 pathways (subgraphs) that connect ES HSCs to CMPs and CLPs (Extended Data Fig. 4a) and translated these into kinetic equations describing cell abundance, RFP label propagation and H2B-GFP dilution (Supplementary methods, Section 1). The resulting pathway models were ranked by their ability to fit the experimental data, using the bias-corrected Akaike information criterion (AICc^28^). Remarkably, a single model was substantially better than the next-ranked models (ΔAICc>2; Fig. 2b), showing that high Sca-1 levels were either inherited from precursor (e.g. Sca-1^hi^ HSC) to progeny (e.g. Sca-1^hi^ MPP) or lost irreversibly (Fig. 2c); this model fit all the experimental data (Fig. 2d-f; for parameter values see Supplementary Table 1). The next best ranking models (within ΔAICc<10) were variants of the best model (indicated by orange bars in Fig. 2b; Extended Data Fig. 4b). Models that either lacked inheritance of Sca-1 level (Fig. 2b, green bars, non-parallel) or allowed Sca-1^lo^ to Sca-1^hi^ transitions (Fig. 2b, blue bars), or both (Fig. 2b, grey bars), did not account for the experimental data.

**Figure 2.**
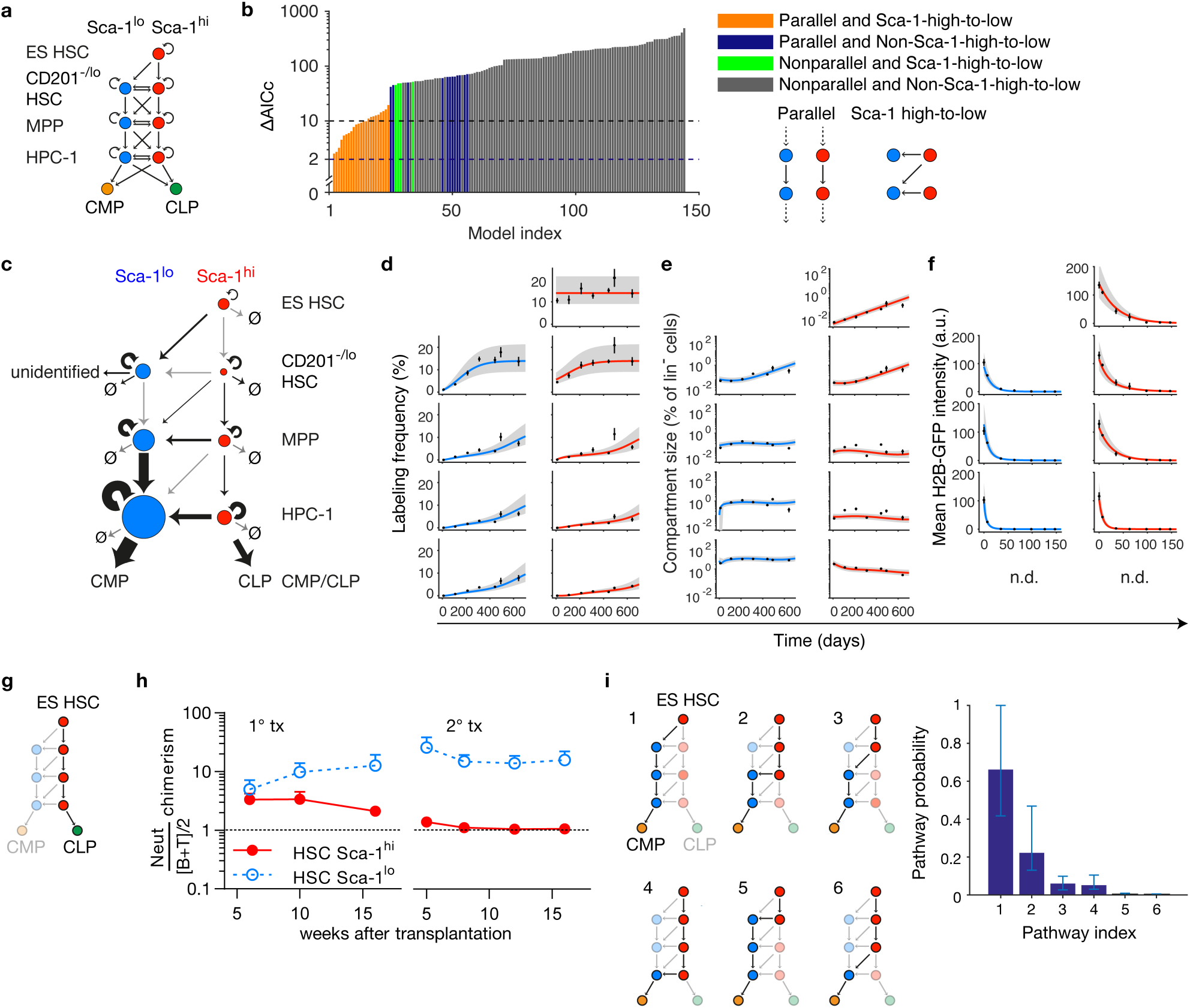
Statistical inference of HSC differentiation pathways. **a**, Model scheme, with subdivision of HSCs and progenitors according to Sca-1 expression and, for HSCs, also CD201 expression. Straight arrows denote cell differentiation (vertical), or change in Sca-1 expression (horizontal), or both (diagonal); curved arrows denote cell proliferation. All potentially possible differentiation pathways emanating from ES HSCs are superimposed. **b**, Model selection by bias-corrected Akaike information criterion (AICc) against the experimental data for HSC fate mapping, mitotic history tracking and relative cell numbers. Models are classified (and color-coded accordingly) by two features: parallel differentiation pathways and irreversible loss of Sca-1. The best-fitting model (shown in **c** below) has ΔAICc = 0 and carries substantially higher evidence than any of the remaining 143 models, as none of these models is within ΔAICc < 2. Further models with acceptable evidence (ΔAICc < 10) are all variants of the best-fitting model (see Extended Data Fig. 4b). **c**, Best model selected by the experimental data. The arrow width signifies the values of the corresponding differentiation or proliferation rates, while the size of a node quantifies the cell number (relative to lin^−^ cells) in each compartment at 100 days of chase; see Supplementary Table 1. **d – f**, Fits of the best model for RFP labeling frequency propagation (**d**), compartment size (**e**) and H2B-GFP dilution (**f**) (experimental data: mean ± SEM, for n = 5 to 12 mice per data point; model fit: straight line, best fit; grey-shaded areas, 95% confidence bounds). **g**, A single pathway connects ES HSCs to CLPs, via Sca-1^hi^ intermediates. **h**, As predicted by the model, Sca-1^hi^ HSCs, but not Sca-1^lo^ HSCs, yielded lymphoid progeny. 100 Sca-1^hi^ or Sca-1^lo^ donor HSCs were competitively transplanted and contribution to peripheral blood neutrophils, B and T cells was determined (data re-analysed from Morcos et al., 2017). Lineage bias of donor-derived leukocytes was calculated (Neutrophil chimerism / [B+T cell chimerism]/2), mean and SEM are shown). **i**, Inferred lineage pathways to CMPs and their relative contributions to generation of CMPs (showing the probability that a CMP was generated by each pathway; error bars, 95% prediction bands).

The selected model (Fig. 2c) makes several predictions: First, it yields a high loss rate of CD201^−/lo^ Sca-1^lo^ HSCs, besides their contribution to Sca-1^lo^ MPPs, implying a contribution to an as yet unidentified downstream population (Fig. 2c, arrow labeled “unidentified”). Second, the model implies a single route from ES HSCs to CLPs, via successive Sca-1^hi^ precursor stages (Fig. 2g). Third, the model predicts that lower Sca-1 expression in HSCs signals erythro-myeloid commitment. The second and third predictions are supported by the outcome of competitive transplantation of Sca-1^hi^ versus Sca-1^lo^ HSCs into lethally irradiated recipient mice (Fig. 2h), while the first prediction will be elucidated below.

In our model, multiple possible routes lead to CMPs, only two of which proceed via CD201^−/lo^ Sca-1^lo^ HSCs (Fig. 2i, pathways 1 and 5). Based on the differentiation rates inferred from the experimental data, we computed the probability that a CMP was generated by any particular pathway (Supplementary methods, Section 2). The majority (~2/3) of CMPs could be traced back to CD201^−/lo^ Sca-1^lo^ HSCs, downstream followed by massive selective proliferation in the Sca-1^lo^ pathway. Hence, in unperturbed hematopoiesis the major commitment to erythro-myeloid differentiation occurs already at the HSC stage.

## CD201^−/lo^ Sca-1^lo^ HSCs feed thrombopoiesis through CD48^−/lo^ MkPs

MPPs have been thought to supply all lineage pathways^13,14^, but recent work indicated that platelets can be generated directly from HSCs in mice^10–12^ and humans^29^. Indeed, when comparing labeling frequencies in committed progenitors and mature cell types relative to MPPs, only platelets showed significantly faster and higher label acquisition than both MPPs and CMPs (Fig. 3a, Extended Data Fig. 5), arguing against thrombopoiesis proceeding solely via MPPs. By contrast, CD201^−/lo^ Sca-1^lo^ HSCs had higher labeling frequency than platelets throughout the entire chase period (Fig. 1e). These data suggest that megakaryocyte progenitors are the unidentified progeny of CD201^−/lo^ Sca-1^lo^ HSCs in the model fit (Fig. 2c), implying two distinct pathways of thrombopoiesis (Fig. 3b). To test this idea, we performed further fate mapping experiments that included MkPs. We found that MkPs had consistently lower RFP labeling frequencies than platelets, which argues against the hypothesis that thrombopoiesis proceeds from HSCs via MkPs to platelets (Fig. 3c). Thus, paradoxically, the fate mapping data rule out thrombopoiesis proceeding either through the conventional pathway via MPPs to MkPs or through the direct pathway from HSCs immediately to a homogeneous MkP population.

**Figure 3.**
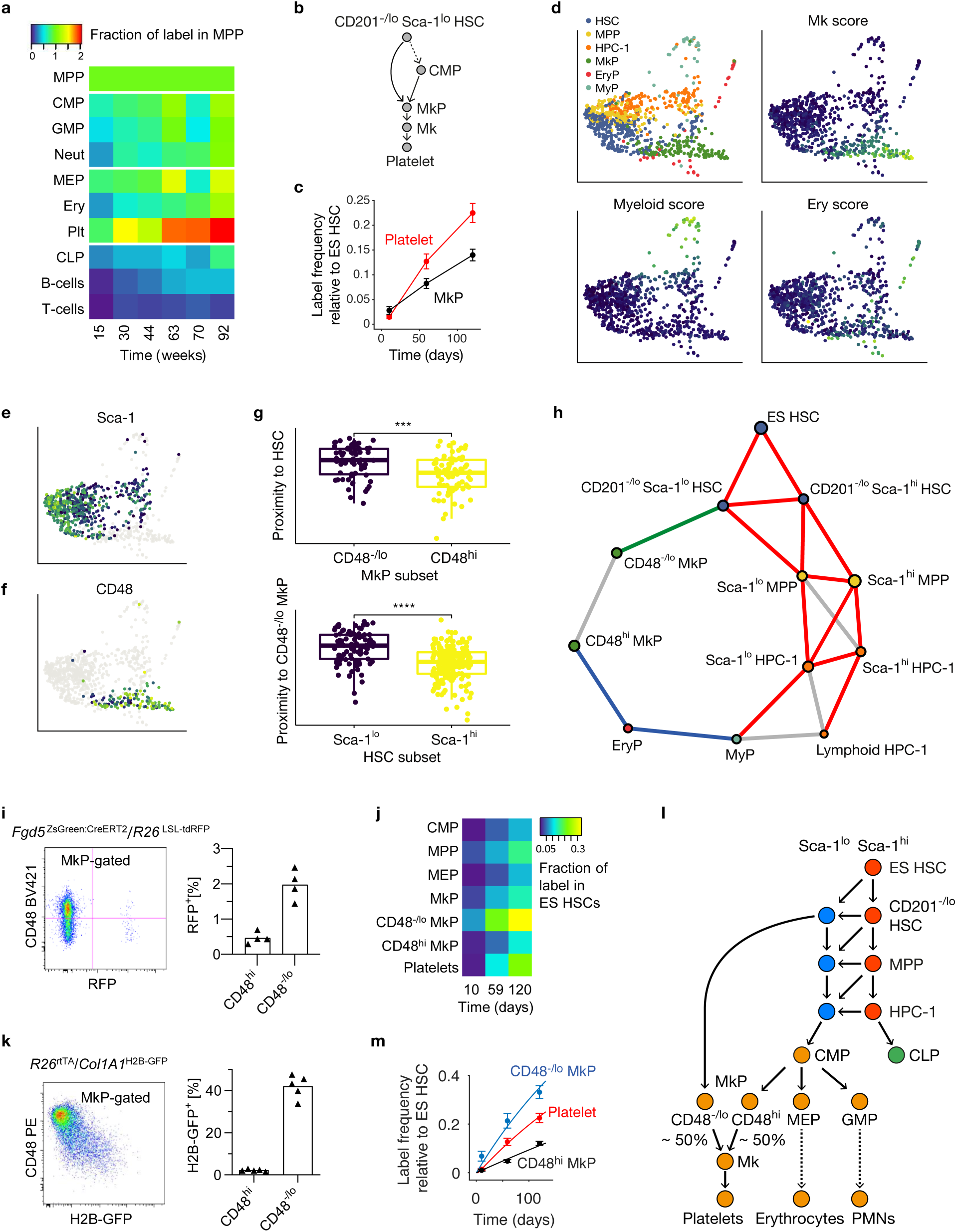
Direct and conventional thrombopoiesis pathways proceed via distinct progenitors. **a**, RFP labelling of cell populations from *Fgd5*^ZsGreen:CreERT2^*/R26*^LSL-tdRFP^ mice (same as in Fig. 1c-e) is shown as fraction of label in the MPP compartment of the same animal at final analysis. **b**, Model scheme with a direct route to MkPs (LS-K CD41^+^CD150^+^) from CD201^−/lo^ Sca-1^lo^ HSCs. **c**, RFP labeling frequency of MkPs is consistently lower than of platelets. **d**, PHATE embedding of single-cell transcriptomes of index-sorted hematopoieitic stem and progenitor cells (n=820 cells in total isolated from 6 *R26*^rtTA^/*Col1A1*^H2B-GFP^ animals, coloured by cell type and expression of lineage markers (blue: low expression, yellow: high expression). **e**, HSCs, MPPs and HPCs-1 show a decrease of Sca-1 surface expression in proximity of MkPs and myeloid-committed progenitors. **f**, Lower surface expression of CD48 in MkPs residing closer to HSCs in the transcriptional landscape. **g**, Proximity in transcription space calculated for MkP subsets to HSCs (top), and for HSC subsets to CD48^−/lo^ MkPs (bottom), placing CD48^−/lo^ MkPs close to Sca-1^lo^ HSCs. **h**, PAGA embedding of surface marker-defined populations, displaying transcriptional proximity of CD201^−/lo^ Sca-1^lo^ HSC to CD48^−/lo^ MkP (green link), and a distinct differentiation path to CD48^hi^ MkP (blue) via myeloid (MyP, LS^−^K CD41C^−^D150^−^) and erythroid progenitors (EryP, LS^−^K CD41^−^CD150^+^). Red PAGA links were previously inferred by statistical model selection against label propagation data (see Figure 2c). Cell populations were identified via indexed surface markers, except for lymphoid-biased HPCs-1, which were identified by their transcriptional signature. **i, j** *Fgd5*^ZsGreen:CreERT2^*/R26*^LSL-tdRFP^ mice were TAM-induced and indicated cell populations were analysed at 10, 59 and 120 days. **i**, Representative example of enriched RFP labeling in CD48^−/lo^ MkPs 59 days after TAM induction. Quantification of label in MkP subpopulations (left data plot, individuals and means are shown). **j**, RFP labeling of cell populations displayed as fraction of ES HSCs. **k**, *R26*^rtTA/rtTA^/*Col1A1*^H2B-GFP/H2B-GFP^ animals (n=5) were DOX-induced and chased for 11 wks. Representative dot plot (left) and frequency of label retaining (H2B-GFP^+^) MkP subpopulations (right, individual mice and means) are shown. **l**, Full model of thrombopoiesis. The direct (via CD48^−/lo^ MkP) and the conventional (via CMP-CD48^hi^ MkP) pathways make equal contributions to native thrombopoiesis (see Extended Data Figure 6h). **m**, Mathematical inference on the fate mapping data (**i,j**) based on the two-pathway model of thrombopoiesis (**l**) fits the experimental data and confirms the prior model prediction that MkPs consist of two distinct subpopulations (see Extended Data Fig. 6b).

As MkPs are well established as precursors of megakaryocytes (Mks)^30,31^, we searched for solutions of this paradox. The simplest working model, as confirmed by mathematical analysis of RFP label propagation, suggests that the conventional and the direct thrombopoiesis pathways are active simultaneously *and* proceed through distinct MkP subpopulations (Extended Data Fig. 6a-b).

To gain unbiased insight into putative differentiation trajectories from HSCs to MkPs, we performed single-cell RNA sequencing (scRNAseq) of index-sorted HSCs, MPPs, HPCs-1 and LS^−^K progenitors including MkPs, using Smart-seq2 ^32^. These populations occupied distinct but partially overlapping regions in the transcriptional landscape computed by PHATE^33^ (Fig. 3d, upper left panel). The different kinds of progenitor cells were well identified by the phenotypic markers used, as indicated by preferential expression of myeloid, erythroid and megakaryocyte lineage gene modules in myeloid progenitors, erythroid progenitors, and MkPs, respectively (Fig. 3d). Moreover, Sca-1 was most highly expressed in HSCs at the apex of the transcriptional landscape (Fig. 3e). We noted that MkPs showed heterogeneous expression of CD48 (Fig. 3f), with CD48^−/lo^ MkPs being significantly closer to CD201^−/lo^ Sca-1^lo^ HSCs in transcript space (Fig. 3g). To extract putative developmental trajectories between HSC and progenitor subpopulations, we applied the graph abstraction method PAGA^34^ (Fig. 3h), which almost completely recovered the model topology inferred from fate mapping (compare Fig. 3h, red lines with Fig. 2c). Remarkably, for thrombopoiesis, only CD48^−lo^ MkPs were directly connected to CD201^−/lo^ Sca-1^lo^ HSCs (Fig. 3h, green line) whereas CD48^hi^ MkPs were connected to HPC-1 via LS^−^K progenitors (Fig. 3h, blue lines). Thus, single-cell transcriptome data point to distinct origins of CD48^−/lo^ and CD48^hi^ MkPs.

PAGA also inferred a putative link between CD48^−/lo^ and CD48^hi^ MkPs, indicating that they could be connected. Importantly, however, while developmental connection implies proximity in transcript space the reverse need not be true. To address this, we performed functional *in vivo* assays using fate mapping. Indeed, fate-mapping label was enriched among CD48^−/lo^ MkPs in recently induced *Fgd5*^ZsGreen:CreERT2^*/R26*^LSL-tdRFP^ mice, whereas CD48^hi^ MkPs acquired less label (Fig. 3i). Continued chase confirmed more rapid and higher label acquisition by CD48^−/lo^ MkPs versus CD48^hi^ MkPs (Fig. 3j). Moreover, the complete lack of label equilibration between the two subsets implies that they are located in two distinct pathways (rather than CD48^−/lo^ MkPs being upstream of CD48^hi^ MkPs). As predicted by PAGA, CD48 MkPs were developmentally close to HSCs, as they were the only cell type among all LS^−^K progenitors which retained or inherited significant amounts of H2B-GFP in chased *R26*^rtTA^/*Col1A1*^H2B-GFP^ mice (Fig. 3k) Further characterization of the functional potential of CD48^−/lo^ or CD48^hi^ MkPs *in vitro* and post transplantation (Extended Data Fig. 6c-g) confirmed their predominant thrombopoietic potential.

To summarize, combining the data from scRNAseq, fate mapping and mitotic tracking, we identified a pathway of thrombopoiesis proceeding directly from CD201^−/lo^ Sca-1^lo^ HSCs to CD48^−/lo^ MkPs that is used in parallel with the conventional pathway of platelet generation via MPPs and CD48^hi^ MkPs (Fig. 3l). Inferring the flux through these pathways (Fig. 3m, Extended Data Fig. 6h), we find that they both contribute to steady-state thrombopoiesis at a similar magnitude (Fig. 3l).

## Thrombopoietin signaling enhances direct HSC-derived thrombopoiesis

The differentiation rate of HSCs is thought to respond to the need for mature cell types after injury or other challenges, but a direct assay for assessing HSC differentiation in physiological hematopoiesis has been lacking. We reasoned that HSC fate mapping may provide such an assay. To examine this, we exposed TAM-induced *Fgd5*^ZsGreen:CreERT2^*/R26*^LSL-tdRFP^ and DOX-pulsed *R26*^rtTA^/*Col1A1*^H2B-GFP^ mice to the myeloablative drug 5-Fluorouracil (5-FU), and investigated how ES HSCs contribute to hematopoietic recovery (Fig. 4a, b). 5-FU substantially accelerated the contribution of RFP-labelled HSCs to progenitors and mature cells (Fig. 4c, Extended Data Fig. 7a), consistent with ^2,7^. Along with increased HSC output, 5-FU provoked H2B-GFP dilution equivalent to ~3-4 additional cell divisions in all HSC and MPP subpopulations (Extended Data Fig. 7b). In more proliferative progenitors (e.g. HPC-1, CD48^hi^ MkP), 5-FU diluted the H2B-GFP label to the range of the background control. By contrast, CD48^−/lo^ MkPs still exhibited H2B-GFP labeling after recovery from 5-FU treatment, which further supports their direct emergence from HSCs. Taken together, these experiments show that HSC fate mapping provides an assay for changes in HSC output.

**Figure 4.**
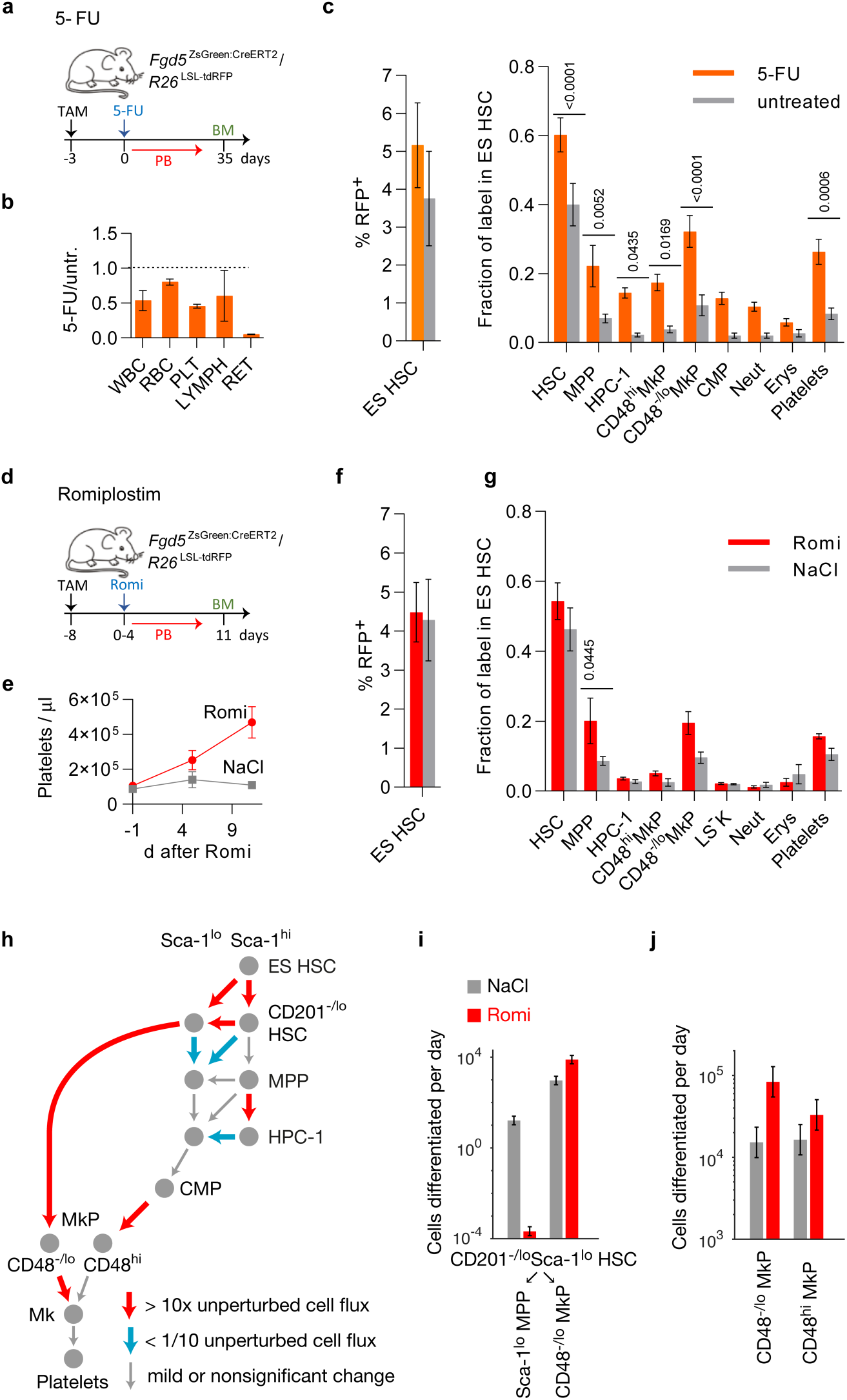
Fate mapping of perturbed hematopoiesis. **a-c**, *Fgd5*^ZsGreen:CreERT2^*/R26*^LSL-tdRFP^ mice were TAM-induced, treated with 5-FU (n=5) or left untreated (n=5), sacrificed at various time points, and bone marrow populations and peripheral blood leukocytes were analysed for RFP expression. **b**, Ratios of peripheral blood cell numbers in treated (5-FU) vs. untreated (untr.) animals 6 days after treatment. **c**, Label in populations 35 days after treatment is shown as percentage of RFP^+^ cells in ES HSC (left graph) or normalised as fraction of label in ES HSC (right graph, mean with SEM, one-way ANOVA with Sidak correction). **d-f**, *Fgd5*^ZsGreen:CreERT2^*/R26*^LSL-tdRFP^ mice were TAM-induced, treated with romiplostim (n=4) or saline (NaCl, n=4), and sacrificed 11 d after start of treatment and hematopoietic populations were analysed for RFP expression. **e**, Time-course of platelet numbers. **f**, RFP labelling in ES HSCs. **g**, Label in bone marrow and blood populations is shown as fraction of label in ES HSC in respective populations (one-way ANOVA with Sidak correction). **h**, Graphical overview of altered cell fluxes (number of cells differentiated per day) by romiplostim application. Red and blue arrows represent up-regulation and down-regulation, respectively, over one order of magnitude. Grey arrows indicate minor or non-significant changes. (cf. Extended Data Fig. 8). **i, j**, Cell fluxes of either CD201^−/lo^ Sca-1^lo^ HSCs into Sca-1^lo^ MPPs and CD48^−/lo^ MkPs (**i**) or via CD48^−/lo^ and CD48^hi^ MkP (**j**) in steady state (NaCl, grey) or after romiplostim (red) perturbation.

We used this assay to study the effects of thrombopoietin signaling, which is a physiological regulator of thrombopoiesis as well as HSC maintenance^35^. To this end, we treated *Fgd5*^ZsGreen:CreERT2^*/R26*^LSL-tdRFP^ and *R26*^rtTA^/*Col1A1*^H2B-GFP^ mice with the thrombopoietin receptor (Mpl) agonist, romiplostim, for 5 consecutive days (Fig. 4d). As previously reported^36^, this regimen expanded all LSK sub-populations except MPPs (Extended Data Figure 7c and raised platelet counts (Fig. 4e). To understand how this is achieved, we studied label propagation from ES HSCs. RFP labeling frequencies of total HSCs including ES HSCs were not affected by romiplostim treatment (Fig. 4f). However, we observed strong increases of RFP-labelling frequencies in downstream progenitors, including MPPs and CD48^−/lo^ MkPs, implying accelerated differentiation of HSCs into these progenitors (Fig. 4g, Extended Data Fig. 7a). In addition, romiplostim administration to *R26*^rtTA/^*Col1A1*^H2B-GFP^ mice induced ~1-2 additional divisions in HSCs and progenitors (Extended Data Fig. 7d). To understand how acceleration of differentiation and enhanced cell division conspire to increase platelet production, we fitted the mathematical model to the experimental data (Fig. 3m, Extended Data Fig. 8a-c). Flux changes affected primarily thrombopoiesis through the direct pathway (Fig. 4h). In particular, CD201^−/lo^ Sca-1^lo^ HSCs became repurposed in a coordinated manner, by decreasing their output to Sca-1^lo^ MPPs and increasing their output to CD48^−/lo^ MkPs (Fig. 4i). In sum, this increased the rate of platelet production through the direct pathway by one order of magnitude whereas the conventional pathway was only moderately upregulated (Fig. 4j). Taken together, thrombopoietin signaling enhances platelet production by channeling CD201^−/lo^ Sca-1^lo^ HSCs into the direct thrombopoiesis pathway.

We have demonstrated that stem cell fate mapping charts physiological differentiation pathways when combined with proliferation assays and mathematical inference. We thus pinpoint the tip stem cells of native, steady-state hematopoiesis, show how lineage pathways diverge immediately downstream of tip HSCs, and find that thrombopoiesis proceeds via two distinct progenitors that previously were thought to be a homogeneous population. Data integration by using computational techniques emerges as a highly promising direction for understanding physiological tissue renewal. Specifically, the identification of hitherto unrecognized distinct MkPs has been enabled by closely linking single-cell transcriptomics with fate mapping. Moreover, the accuracy of predictions derived from computational model selection attest to the utility of this tool for interrogating complex, multi-faceted data. Surface markers of stem and progenitor cells have been selected for their utility in prospective isolation of these cells, and much less emphasis has been given to their molecular function. Interestingly, however, Sca-1, marking tip-HSCs together with CD201, is an interferon-stimulated gene (ISG)^37^. Constitutive expression of ISGs, with protective function, is a hallmark of stem cells also beyond the hematopoietic system^38^. CD201 has been implied in HSC retention in the niche^39^. Our functional data on tip HSCs imply that the gene expression program associated with high co-expression of Sca-1 and CD201 deserves closer scrutiny.

Our identification of a completely separate pathway of thrombopoiesis, leading from Sca-1^lo^ HSC via CD48^−/lo^ MkPs to megakaryocytes, bolsters the recently advanced view that thrombopoiesis is special among all adult hematopoietic lineage pathways by emerging directly from HSCs already under steady-state conditions^10,11,29^. Contrary to the classical view^13,17^, but in agreement with^10^, this pathway does not proceed via MPP. Our quantification of the cell fluxes from HSC subpopulations to progenitors now also suggest a functional rationale for this seemingly peculiar feature of mammalian hematopoiesis. While platelets, neutrophils and erythrocytes make up the major share of hematopoietic cell production in steady state^40^, their amplification strategies are fundamentally different: Neutrophils and erythrocyte numbers are achieved by massive progenitor proliferation, whereas platelet numbers are obtained via prolonged shedding from megakaryocytes after enormous cell growth. We now find that this dichotomy is already manifested at the HSC level. The major cell flux from Sca-1^lo^ HSCs supplies thrombopoiesis, while myelopoiesis is fed by a much smaller flux from Sca-1^lo^ HSCs and amplified by selective proliferation of Sca-1^lo^ progenitors downstream. Our observation that activating a physiological regulator of thrombopoiesis further accentuates this effect (increasing the flux from Sca-1^lo^ HSCs to thrombopoiesis and accelerating, in a compensatory manner, proliferation of Sca-1^lo^ MPPs) strengthens this interpretation. Thus, our data show that HSCs contribute very actively to thrombopoiesis in native hematopoiesis, whereas they feed only infrequently into myelopoiesis, which is sustained primarily by progenitor amplification. Accordingly, mice with a severe reduction of HSCs feature normal numbers of leukocyte and erythrocytes but a significant reduction of platelets^41^.

In sum, the approach developed here connects the charting of lineage pathways with the quantification of cell fluxes. It therefore has great potential for the quantitative understanding of the regulation of native hematopoiesis.

## Acknowledgments

T.H. was supported by the German Research Council (DFG) through SFB 873 (B11) and SFB 1129 (B11), and European Union project 764698-Quantitative T-Cell Immunology and Immunotherapy (QuanTII). A. Gerbaulet was supported by DFG (GE3038/1-1) and Fritz Thyssen Stiftung. A.R. and A.D. were funded by DFG (RO2133/10-1) in the setting of FOR2577. M.N.F.M. was supported by the Dresden International Graduate School for Biomedicine and Bioengineering, granted by DFG in the context of the Excellence Initiative. We thank Christa Haase, Madeleine Rickauer and Livia Schulze for technical assistance and all members of Höfer and Gerbaulet groups for discussion.

## Methods

### Mice

All animal experiments were conducted according to institutional guidelines and in accordance with the German Law for Protection of Animals approved by Landesdirektion Dresden (TVV 91/2017). Mice were housed in individually ventilated cages under specific-pathogen free environment at the Experimental Center of the Medical Faculty, TU Dresden. *Fgd5*^ZsGreen:CreERT2/wt^*/R26*^LSL-tdRFP/LSL-tdRFP 1, 19^, *R26*^rtTA/rtTA^ */Col1A1*^H2B-GFP/H2B-GFP^ (Jax No: 016836) ^42^, C57Bl/6JRj wt (Janvier), B6.CD45.1 (Jax No. 002014) and B6.CD45.1/.2 mice were used in this study. B6.RFP mice with ubiquitious tdRFP expression were generated by germline excision of the lox*P* flanked STOP cassette (LSL) in *R26*^LSL-tdRFP^ animals employing the pgk-Cre transgene^43^. Pulse-chase data of unperturbed *R26*^rtTA^*/Col1A1*^H2B-GFP^ mice was previously published^27^. H2B-GFP mice were induced with doxycyclin (DOX) via chow (Ssniff Spezialdiäten, 625mg/kg for 10-11 wks or 2000 mg/kg for either 2-7 wks (this study) ad libitum. *Fgd5*^ZsGreen:CreERT2^*/R26*^LSL-tdRFP^ mice were induced by oral gavage of tamoxifen (0.2 mg/g BW) twice 3-4 days apart. The Mpl agonist romiplostim (Nplate, Amgen, 2.5 µg/mouse/d) was injected i.p. for 5 consecutive days. 5-FU (150 µg/g body weight, Applichem) was administered via intravenous (i.v.) injection.

### Cell Preparation

Whole bone marrow cells were isolated by crushing long bones with mortar and pestle using PBS/2% FCS/2 mM EDTA and filtered through a 100 µm mesh. After erythrocyte lysis in hypotonic NH4Cl-buffer, cells were filtered through a 40 µm mesh. Hematopoietic lineage^+^ cells were removed with the lineage cell depletion kit (Miltenyi Biotec).

Peripheral blood was drawn into glass capillaries by retro-bulbar puncture. For identification of RFP^+^ platelets and erythrocytes, 1-2 µl of heparinized blood was mixed with PBS/2%FCS/2mM EDTA and incubated with monoclonal antibodies against CD41 and Ter119 (for gating see Extended Data Fig. 1d). For leukocyte analysis, erythrocyte lysis in hypotonic NH4Cl-buffer was performed twice for 5 min and cells were stained with monoclonal antibodies (Extended Data Fig. 1d). For hemograms, blood was drawn by retro-bulbar puncture directly into EDTA-coated tubes (Sarstedt) and analyzed on a XT-2000i Vet analyzer (Sysmex).

### Transplantation

B6.CD45.1/.2 recipient mice received a single dose of 9 Gray total body irradiation (Yxlon Maxi Shot γ-source). Donor cells (10 purified LSK CD48^−/lo^ CD150^+^ Fgd5-ZsGreen^+^ test cells mixed with 200000 B6.CD45.1 competitor bone marrow cells) were administered via intravenous injection into the retro-orbital sinus. Peripheral blood T-lymphocytes (CD3^+^), B-lymphocytes (B220^+^) and neutrophils (CD11b^+^, Gr-1^hi^) were analyzed for their donor origin using an Aria III flow cytometer. The frequency of HSCs with long-term multi-lineage repopulation potential was estimated by ELDA^44^. Competitive transplantation of either Sca-1^hi^ or Sca-1^lo^ donor HSCs was previously described^25^. Briefly, 100 test donor HSCs (LSK CD48^/lo^ CD150^+^) from WT B6 mice were sorted for Sca-1 expression level, mixed with 500,000 B6.CD45.1 competitor bone marrow cells and i.v. injected into irradiated B6.CD45.1/.2 recipient mice.

For transplantation of MkP populations, 1000 either CD48^−/lo^ or CD48^hi^ MkPs were purified from B6.RFP donor mice, mixed with 500,000 B6.CD45.1/.2 carrier splenocytes and injected into B6.CD45.1/.2 recipient mice that previously received 4 Gy γ-irradiation. Recipient mice that were transplanted with lin^−^CD201^+^ CD117^+^ (LEK) cells served as positive controls, while animals that received only carrier splenocytes were used as background controls.

### Flow cytometry

Cell suspension were incubated with antibodies (Supplementary Table 2) in PBS/2% FCS/2mM EDTA for 30 min, washed twice and analyzed on either FACS Canto, ARIA II SORP, ARIA III (all from BD Biosciences, Heidelberg, Germany) or MACSquant (Miltenyi) flow cytometers. Data were analyzed with FlowJo V9.9 and V10 software (Tree Star) and gates were set with the help of Fluorescence-Minus-One controls. For a detailed overview of gating strategies refer to Extended Data Fig. 1.

### Single cell culture

Single cells were deposited into 96 well U-bottom plate (TPP) using a BD FACS ARIA II cell sorter. Cells were cultivated in Iscove’s Modified Dulbecco’s Medium (Gibco) supplemented with 10% FCS (FBS superior, Biochrom), 1% Pen/Strep, stem cell factor (rmSCF 20ng/ml, PeproTech), Thrombopoietin (rmTPO 20ng/ml, PeproTech), Interleukin-3 (rmIL-3 20ng/ml, PeproTech), Erythropoietin (rhEPO, 5U/ml, NeoRecormon, Roche). Culture wells were visually inspected and counted by light microscopy (PrimoVert, Zeiss) and assigned to categories on day 5 of culture (see Extended Data Fig. 6 c-d). Representative wells were stained with Hoechst 33342 (5 µg/ml) and imaged by fluorescence microscopy (BZ-X710, Keyence).

### Single-cell RNA sequencing

Single hematopoietic stem and progenitor cells were index-sorted into 384 well plates containing 0.5 µl of nuclease free water with 0.2% Triton-X 100 and 4 U murine RNase Inhibitor (NEB), spun down and frozen at −80°C. After thawing, 0.5 µl of a primer mix were added (5 mM dNTP (Invitrogen), 0.5 µM dT-primer (C6-aminolinker-AAGCAGTGGTATCAACGCAGAGTCGACTTTTTTTTTTTTTTTTTTTTTTTTTTTTTTVN), 1 U RNase Inhibitor (NEB)). RNA was denatured for 3 minutes at 72°C and the reverse transcription (RT) was performed at 42°C for 90 min after filling up to 10 µl with RT buffer mix for a final concentration of 1x superscript II buffer (Invitrogen), 1 M betaine, 5 mM DTT, 6 mM MgCl2, 1 µM TSO-primer (AAGCAGTGGTATCAACGCAGAGTACATrGrGrG), 9 U RNase Inhibitor and 90 U Superscript II. After synthesis, the reverse transcriptase was inactivated at 70°C for 15 min. The cDNA was amplified using Kapa HiFi HotStart Readymix (Peqlab) at a final 1x concentration and 0.1 µM UP-primer (AAGCAGTGGTATCAACGCAGAGT) under following cycling conditions: initial denaturation at 98°C for 3 min, 23 cycles [98°C 20 sec, 67°C 15 sec, 72°C 6 min] and final elongation at 72°C for 5 min. The amplified cDNA was purified using 1x volume of hydrophobic Sera-Mag SpeedBeads (GE Healthcare) resuspended in a buffer consisting of 10 mM Tris, 20 mM EDTA, 18.5 % (w/v) PEG 8000 and 2 M sodium chloride solution. The cDNA was eluted in 12 µl nuclease free water and the concentration of the samples was measured with a Tecan plate reader Infinite 200 pro in 384 well black flat bottom low volume plates (Corning) using AccuBlue Broad range chemistry (Biotium).

For library preparation up to 700 pg cDNA was desiccated and rehydrated in 1 µl Tagmentation mix (1x TruePrep Tagment Buffer L, 0.1 µl TruePrep Tagment Enzyme V50; from TruePrep DNA Library Prep Kit V2 for Illumina; Vazyme) and tagmented at 55°C for 5 min. Subsequently, Illumina indices are added during PCR (72°C 3 min, 98°C 30 sec, 13 cycles [98°C 10 sec, 63°C 20 sec, 72°C 1 min], 72°C 5 min) with 1x concentrated KAPA Hifi HotStart Ready Mix and 300 nM dual indexing primers. After PCR, libraries are quantified with AccuBlue Broad range chemistry, equimolarly pooled and purified twice with 1x volume Sera-Mag SpeedBeads. This was followed by Illumina 50 bp paired-end sequencing on a Novaseq6000 aiming at an average sequencing depth of 0.5 mio reads per cell.

### Single-cell transcriptome analysis

Raw reads were mapped to the mouse genome (mm10) and splice-site information from Ensembl release 87 ^45^ with gsnap (version 2018–07-04) ^46^. Uniquely mapped reads and gene annotations from Ensembl were used as input for featureCounts (version 1.6.2 ^47^) to create counts per gene and cell. Outlier cells in quality control metrics (library size, number of detected genes, ERCC and mitochondrial reads) were excluded. One out of three plates exhibited a large number of poor quality cells, thus quality control metrics from the remaining two plates were used for outlier detection^48^.Normalization was performed using the pooling method^48^ combined with scaling normalization within each plate to account for different coverages (multiBatchNorm function in the R batchelor package). Batch effects were regressed out of the data using a linear model (removeBatchEffect method in the R limma package). Mutual nearest neighbour correction was also tested, confirming the results from the linear model. Feature selection was performed using variance modelling based on spike-in transcripts and selecting genes with a non-zero biological variance with an FDR threshold of 0.05, then excluding genes contained in the cell cycle entry in Gene Ontology. 15 principal components were retained and used to compute PHATE embedding^33^ and transcriptomic distance. A k-nearest-neighbour graph using ten principal components and k = 15 was used to compute the PAGA connectivities^34^, thus embedded using the Fruchterman-Reingold layout with a threshold of 0.9.

### Mathematical modeling

We use ordinary differentiation equations to describe the dynamics of label propagation, compartment sizes and H2B-GFP dilution (for details see Supplementary methods). Computational inference was performed using the trust-region method to search for the local minimum of the negative log-likelihood. To find the global minimum, the initial parameter values were chosen by Latin hypercube sampling (5000 samples for each model). The tasks above were performed using Data2Ddynamic (D2D) framework, a Matlab-based open source package^49^.

---

**Extended Data Figure 1.**
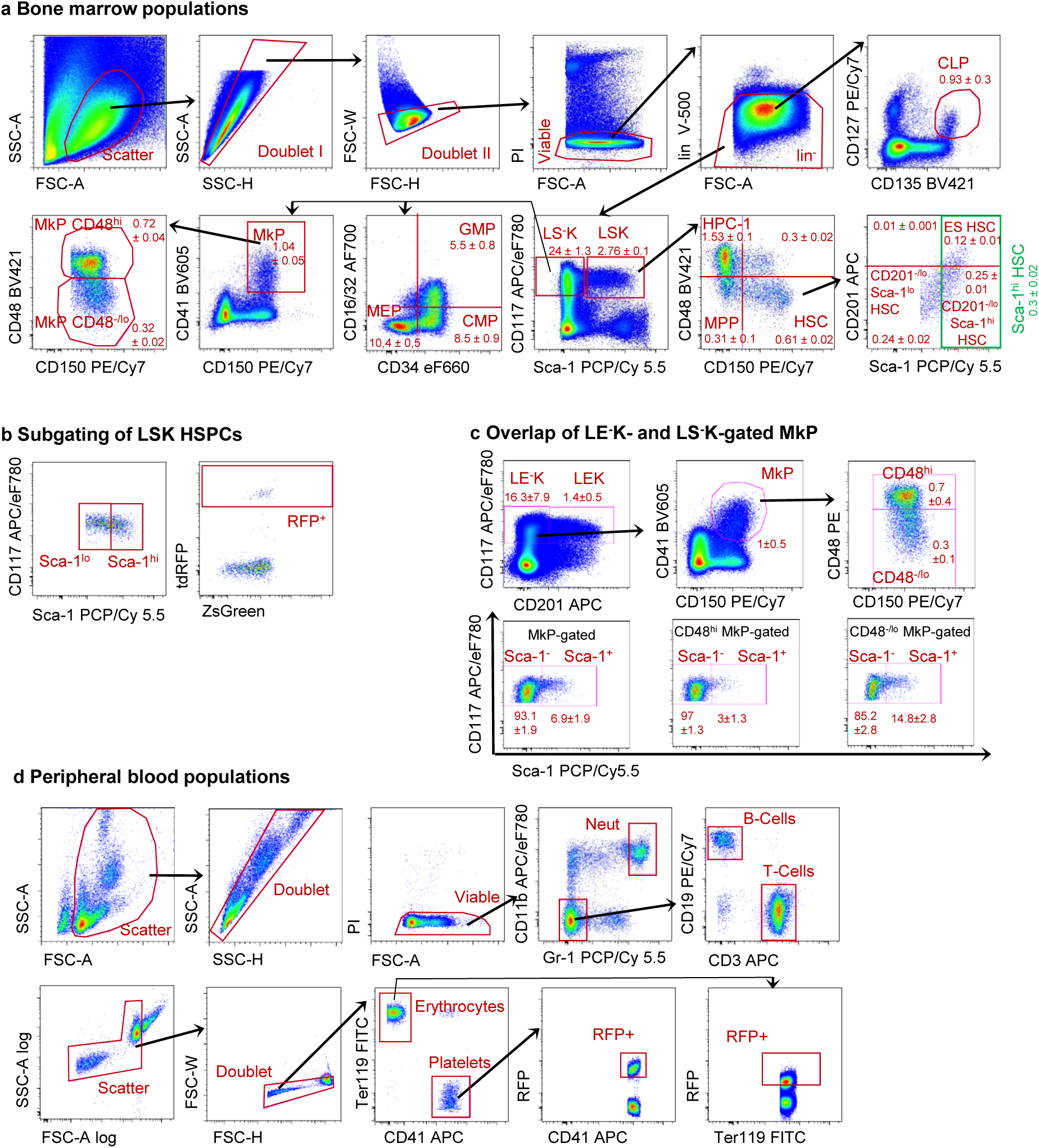
Identification of bone marrow and peripheral blood cell populations. **a**, Representative example of flow cytometry gating for bone marrow HSCs and progenitors; frequencies of populations ± SEM (n=4) among lineage negative cells (lin^−^) are shown (*LS^−^K*, lin^−^ Sca-1^−^ CD117^+^ cells; *LSK*, lin^−^ Sca-1^+^ CD117^+^ cells; *common lymphoid progenitor (CLP)*, lin^−^ CD135^+^ CD127^+^; *megakaryocyte progenitor (MkP)*, LS^−^K CD41^+^ CD150^+^; *granulocyte-macrophage progenitor (GMP)*, LS^−^K CD16/32^+^ CD34^+^; *common myeloid progenitor (CMP)*, LS^−^K CD16/32^−^ CD34^+^; *megakaryocyte-erythrocyte progenitor (MEP)*, LS^−^K CD16/32^−^ CD34^−^; *restricted hematopoietic progenitor 1 (HPC-1)*, LSK CD48^hi^ CD150^−^; *multipotent progenitor MPP)*, LSK CD48^−/lo^ CD150^−^; *hematopoietic stem cell (HSC)*, LSK CD48^−/lo^ CD150^+^; *ES HSC*, CD201^hi^ Sca-1^hi^ LSK CD48^−/lo^ CD150^+^; *CD201*^*−/lo*^ *Sca-1*^*hi*^ *HSC*, CD201^−/lo^ Sca-1^hi^ LSK CD48^−/lo^ CD150^+^; *Sca-1^hi^ HSC*, Sca-1^hi^ LSK CD48^−/lo^ CD150^+^; *CD201*^*−/lo*^ *Sca-1*^*lo*^ *HSC*, CD201^−/lo^ Sca-1^lo^ LSK CD48^−/lo^ CD150^+^)**. b**, Representative sub-gating of LSK populations according to Sca-1 expression level (left). Representative gating of RFP^+^ bone marrow populations (right). **c**, Overlap of lin^−^ CD201^−^ CD117^+^ (LE^−^K) MkPs with the LS^−^K MkP population. **d**, Representative gating of leukocytes (upper row, *neutrophilic granulocytes (Neut)*, CD11b^+^ Gr-1^+^; *B-Cells*, CD11b^−^ Gr-1^−^ CD19^+^; *T-Cells*, CD11b^−^ Gr-1^−^ CD3^+^) and erythrocytes and platelets (lower row, *erythrocytes*, CD41^−^ Ter119^+^; *platelets*, CD41^+^ Ter119^−^).

**Extended Data Figure 2.**
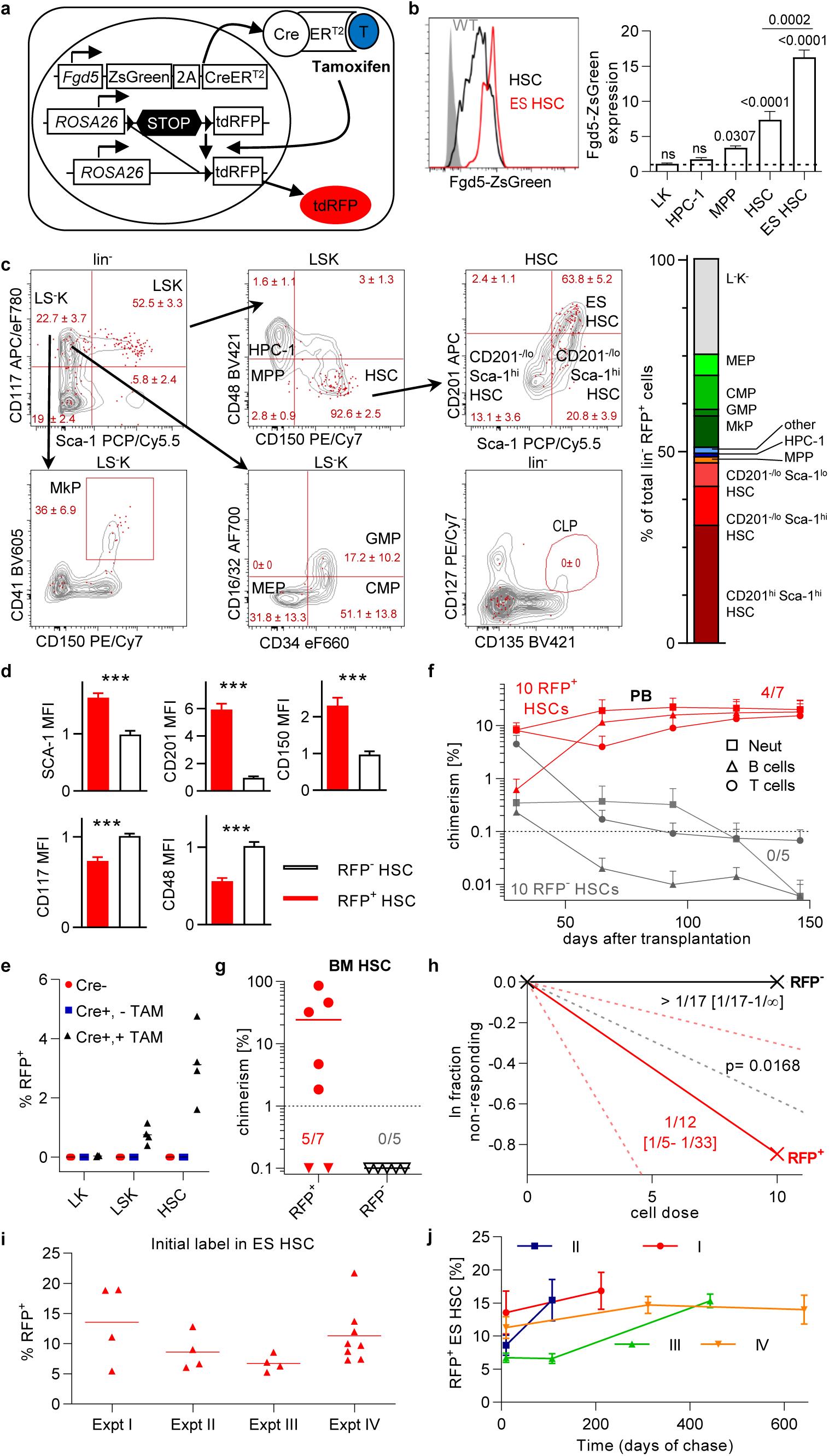
*Fgd5*^ZsGreen:CreERT2^*/R26*^LSL-tdRFP^ mouse model for selective labeling of HSCs. **a**, Schematic of the *Fgd5*^ZsGreen:CreERT2^*/R26*^LSL-tdRFP^ mouse model. The endogenous *Fgd5* promoter drives expression of a green fluorescent protein (ZsGreen) and tamoxifen-inducible Cre-estrogen receptor fusion protein (CreER^T2^). TAM induction causes irreversible excision of the lox*P*-flanked stop cassette (LSL) from the *R26*^LSL-tdRFP^ allele. Cre-recombined cells and their progeny are labeled by stable RFP expression. **b**, ES HSCs (representative histogram shown in red) from *Fgd5*^ZsGreen:CreERT2^*R26*^LSL-tdRFP^ animals express higher levels of Fgd5-ZsGreen than the total HSC (black histogram, autofluorescence control shown in grey) population (paired Student’s t test). Median Fgd5-ZsGreen fluorescence of bone marrow populations isolated from *Fgd5*^ZsGreen:CreERT2^*/R26*^LSL-tdRFP^ animals (n=6, mean ± SEM) was normalized to the autofluorescence of the respective population (dotted line, *Fgd5*^wt/wt^*/R26*^LSL-tdRFP^ mice, n=3). Significant Fgd5-ZsGreen expression was determined by an unpaired one-way ANOVA with Sidak error correction (ns, not significant; p values shown in graph). **c**, *Fgd5*^ZsGreen:CreERT2^*/R26*^LSL-tdRFP^ mice (n=4) were TAM-induced and bone marrow was analysed 10 days later. RFP^+^ lin^−^ cells (red dots) were overlaid to total lin^−^ cells (grey contour plots, representative animal; % ± SEM of RFP^+^ cells within each parental population is shown). RFP^+^ and total bone marrow populations were gated and displayed accordingly. The distribution of hematopoietic populations within total RFP^+^ lin^−^ bone marrow cells is shown (right diagram). Sporadic RFP^+^ cells were detected in the abundant LS-K and L-K-populations and likely reflect slow but continuous differentiation of HSCs within the first 10 days after TAM-induction. **d**, *Fgd5*^ZsGreen:CreERT2^*/R26*^LSL-tdRFP^ mice (n=8) were TAM-induced and bone marrow was analysed 10 days later. Expression (MFI, mean + SEM) of cell surface antigen of RFP^+^ (red bars) and RFP^−^ (open bars) HSCs (LSK CD48^−/lo^ CD150^+^) was normalized to the mean MFI of the total HSC population (significance was calculated by a paired Student’s t test). **e**, *Fgd5*^ZsGreen:CreERT2^*/R26*^LSL-tdRFP^ mice (n=3, age 52 - 57 weeks, Cre^+^, -TAM) were left un-induced and analysed for potential leaky RFP activation. TAM-induced *Fgd5*^ZsGreen:CreERT2^*/R26*^LSL-tdRFP^ (n=4, Cre^+^, +TAM) and Cre-negative (n=4, Cre-) mice served as positive and negative controls, respectively. **f-h**, *Fgd5*^ZsGreen:CreERT2^*/R26*^LSL-tdRFP^ mice were TAM induced and 11 days later either 10 RFP^−^ (grey) or RFP^+^ (red) HSCs (LSK CD48^−/lo^ CD150^+^ ZsGreen^+^) were competitively transplanted into irradiated recipient mice (n = 5 and 7, respectively). **f**, Peripheral blood chimerism (mean and SEM are shown) was analysed and fractions of recipients with successful long-term multi-lineage reconstitution (< 0.1% chimerism (dotted line) in each lineage 146 days after transplantation) are shown. **g**, Bone marrow populations were analyzed 147 days after transplantation for donor-derived cells (individual recipients and means are shown, mice without detectable chimerism are represented by triangles). Fractions of reconstituted mice (>1% chimerism in HSCs (dotted line)) are given. **h**, Frequencies of HSCs with long term (146 days after transplantation) multi-lineage (> 0.1 % in neutrophils, B and T cells, respectively) repopulation activity among transplanted RFP^−^ (shown in black) or RFP^+^ (red) HSCs were estimated by limiting dilution analysis. Frequencies, 95% confidence interval (dotted lines and square brackets) and significance were calculated by ELDA (Hu and Smyth 2009). **i**, *Fgd5*^ZsGreen:CreERT2^*/R26*^LSL-tdRFP^ mice (n=20, same mice as Fig. 1b) were TAM-induced (4 independent experiments (Expt I-IV)) and ES HSCs were analyzed 10 - 11 days after TAM induction for RFP expression. **j**, *Fgd5*^ZsGreen:CreERT2^*/R26*^LSL-tdRFP^ mice (n=74, same mice as in Fig. 1c-e) were TAM-induced (4 independent experiments I-IV) and ES HSCs were analyzed 10 - 642 days after TAM induction for RFP expression. The time course of RFP labeling in ES HSCs for independent experiments is shown.

**Extended Data Figure 3.**
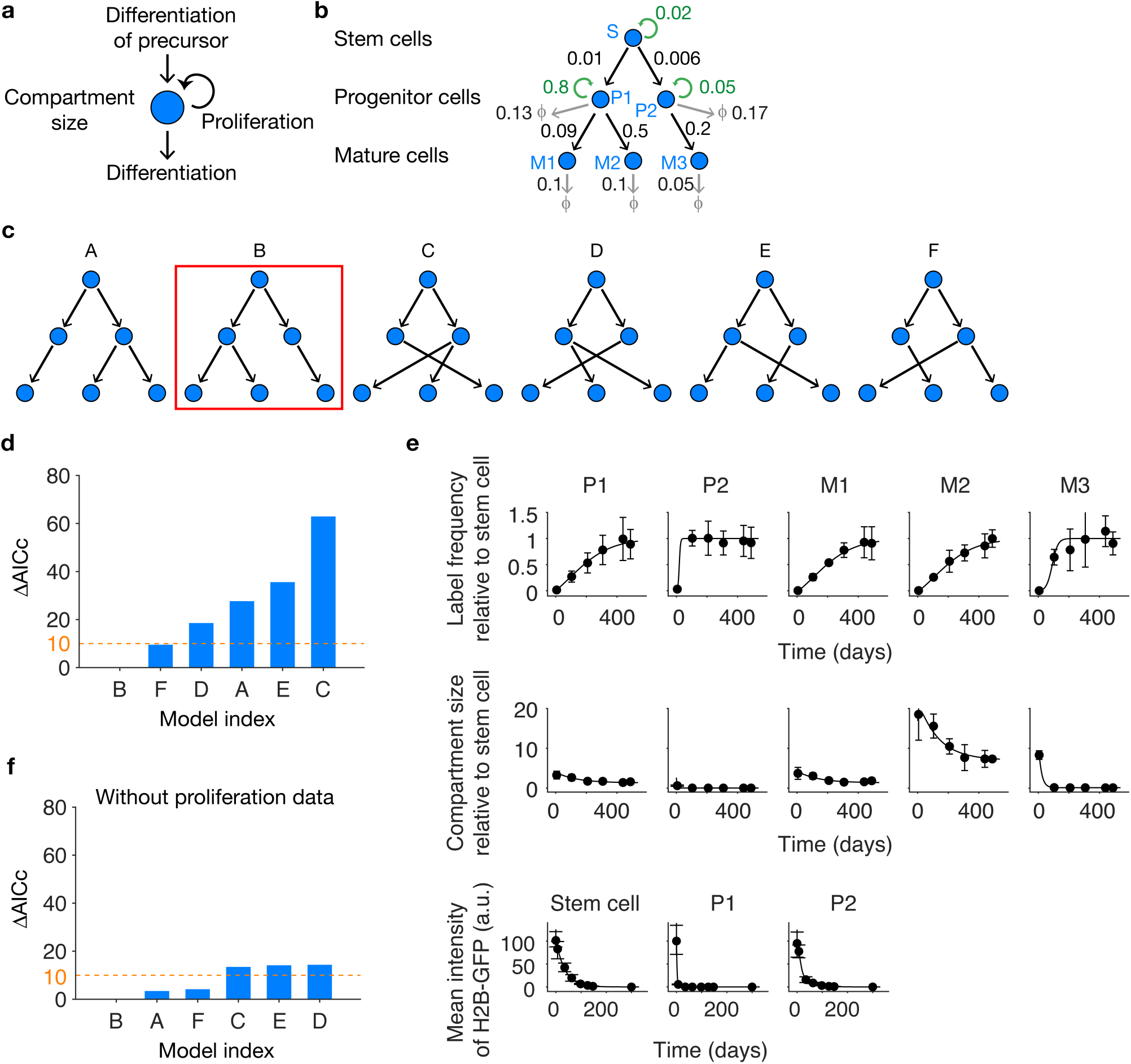
Combining data on HSC fate-mapping and mitotic history allows inference of lineage pathways. **a**, The number of cells in a given population is controlled by the rates of cell differentiation of this population and its precursors and the rate of cell proliferation of this population; in addition, cell death may also occur (not shown). **b**, A generic lineage tree with a stem cell population (S) giving rise to two progenitors (P1 and P2), with P1 yielding mature cell types M1 and M2, and P2 yielding mature cell type M3. To simulate data on cell numbers, fate mapping from S and mitotic history (via H2B-GFP dilution), reasonable values are assigned to cell differentiation rates (black arrows), proliferation rates (green arrows) and death rates (grey arrows), with unit day^−1^. **c**, All possible topologies of lineage pathways connecting S to P1 and P2, and further to M1 through M3. For simplicity, convergence (P1 and P2 spawning the same mature cell type) has not been considered. **d**, Model selection by evaluating how well the schemes A-F in **c** can account for the simulated fate-mapping, H2B-GFP dilution and cell number data computed with the model B. Ranking via the bias-corrected Akaike information criterion (AICc) shows that the original model (B) accounts significantly better for the data than any other model (ΔAICc of the remaining models all > 2). **e**, The best fit of model B (curves) to the simulated data (dots with error bars). **f**, Model selection using only the simulated fate-mapping data.

**Extended Data Figure 4.**
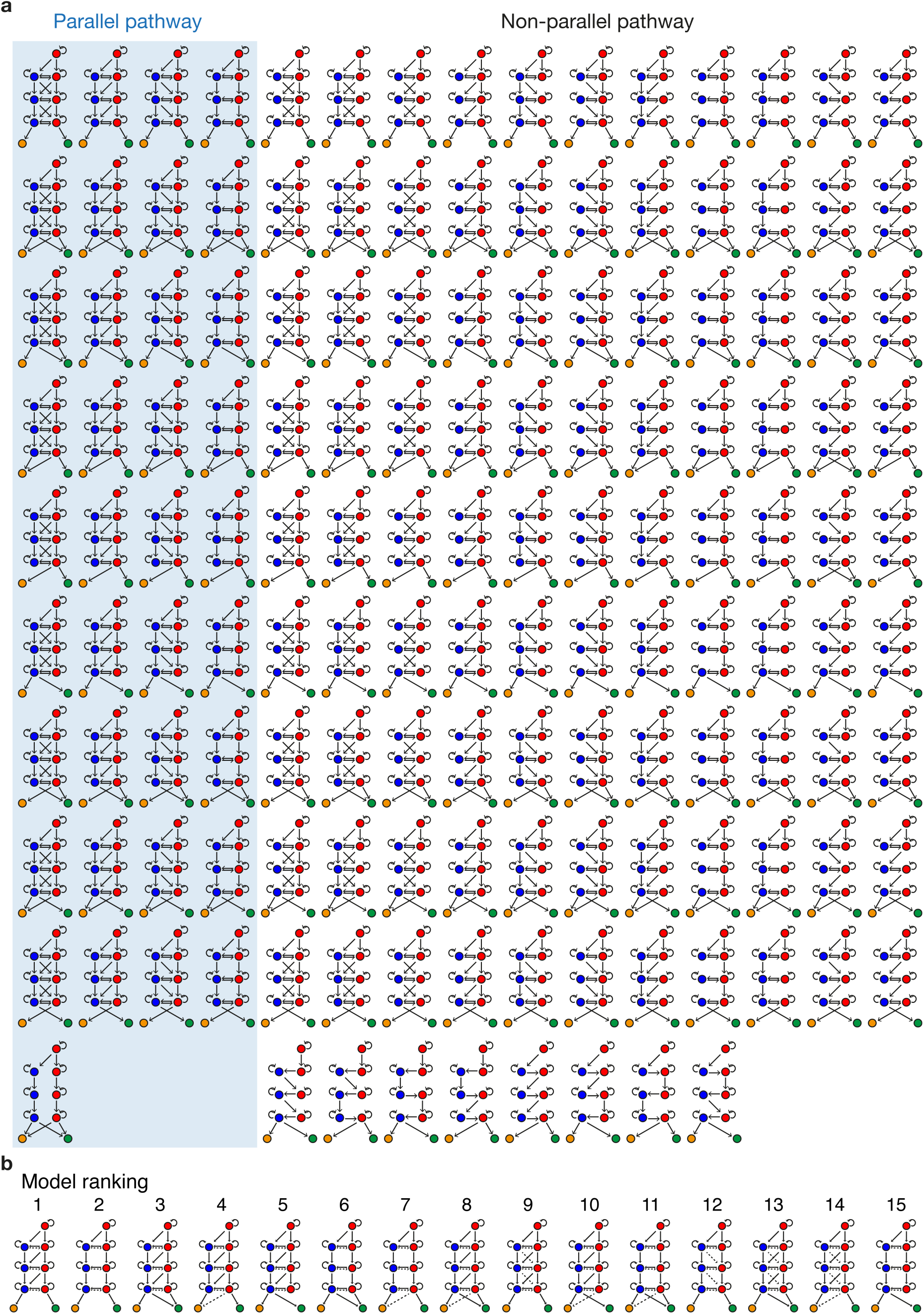
Families of model schemes with different topologies of lineage pathways. **a**, 144 models for statistical model selection. Schemes with parallel Sca-1^hi^ and Sca-1^lo^ pathways are shaded in blue. **b**, Top ranked models with ΔAICc < 10 (dashed arrows, differentiation rate can be 0, as judged by lower 95% confidence bound).

**Extended Data Figure 5.**
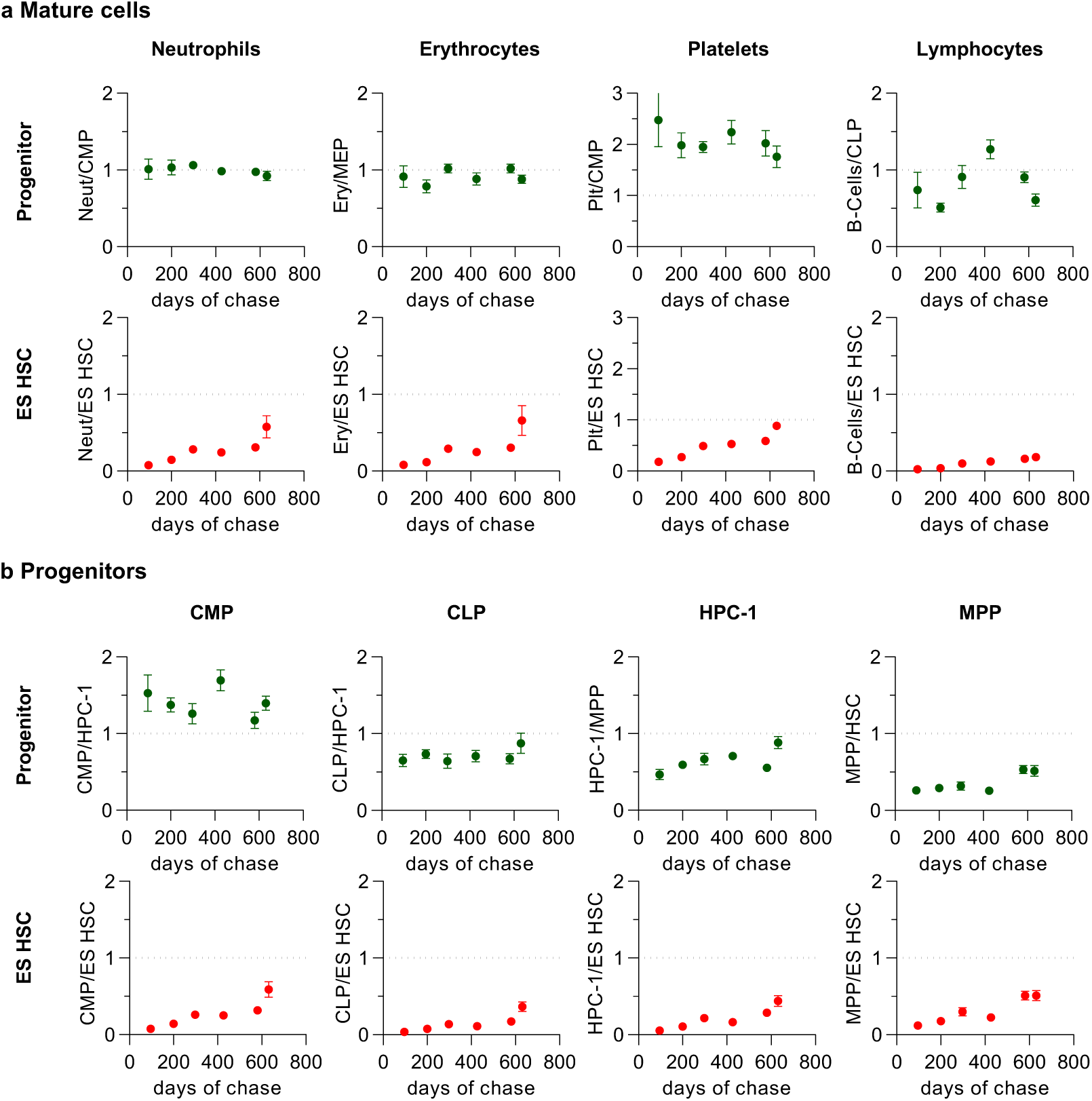
Labeling ratios of HSCs, progenitors and mature cells. **a**, RFP labeling ratios of mature cells from peripheral blood to ES HSCs and their putative progenitors in the bone marrow. Platelets are labeled about twice as frequently as CMPs, indicating that platelets may develop independently of CMPs. **b**, RFP labeling ratios of bone marrow progenitors and ES HSCs. The ratios of RFP labeling were calculated from *Fgd5*^ZsGreen:CreERT2^*/R26*^LSL-tdRFP^ mice (n=82 mice, mean ± SEM, same animals as in Fig. 1c-e).

**Extended Data Figure 6.**
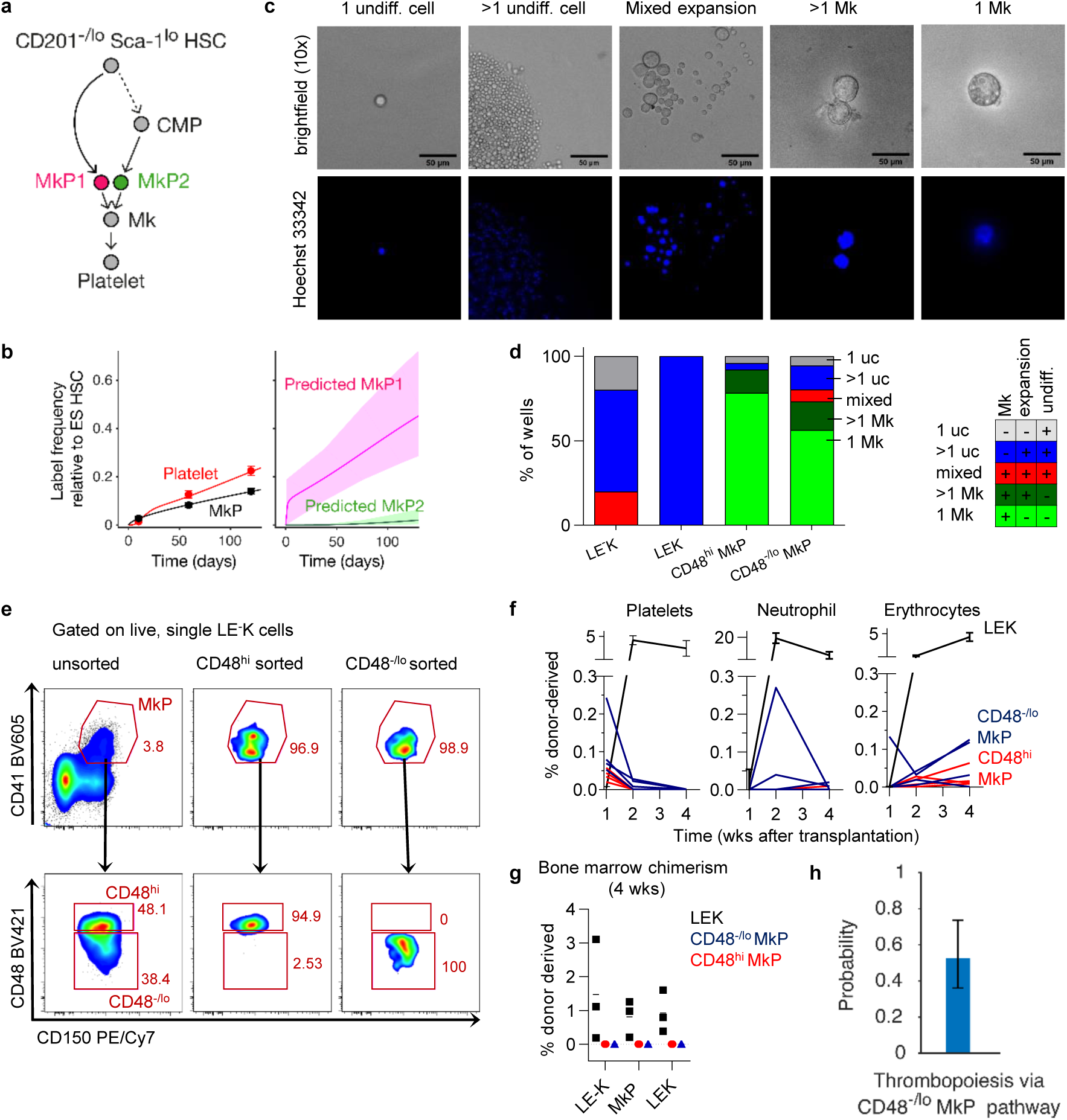
Fate mapping reveals two developmentally distinct MkP subpopulations with similar functional potential. **a**, Hypothetical two-pathway model for MkPs (top), with subpopulations MkP1 (pink) and MkP2 (green). **b**, The two-pathway model fits experimental fate mapping data of platelets and MkPs (left), and predicts distinct labeling frequencies of both MkP subpopulations (means and 95% confidence bounds; right) **c**, Representative examples of single cell cultures inspected by light microscopy (upper row) for cell numbers and morphology. Nuclei were stained by adding Hoechst 33342 (5 µg/ml) and evaluated by fluorescence microscopy (lower row). Growth behavior was assigned to 5 categories: 1 undifferentiated cell (1 uc): seeded cell did neither proliferate nor increase in size; >1 uc: seeded cell divided at least once, but daughter cells did not enlarge; mixed expansion: seeded cell divided at least once and daughter cells were heterogeneous in size; >1Mk: seeded cell divided at least once and all daughter cells enlarged and displayed megakaryocytic morphology; 1 Mk: seeded cells increased in size without dividing. **d**, Single CD48^−/lo^ (n=71) and CD48^hi^ (n=74) MkPs as well as control cells (LE-K: lin^−^ CD201^−^ CD117^+^ progenitors (n=5); LEK: lin^−^ CD201^+^ CD117^+^ cells (n=4)) were cultivated for 5 days and classified according to expansion and morphology. **e**, Purification and re-analysis of CD48^hi^ and CD48^−/lo^ donor MkPs. Frequencies among respective parent populations are shown **f-g**, 1000 CD48^hi^ or CD48^−/lo^ MkPs were purified from B6.RFP donor animals and transplanted into sublethally-irradiated B6.CD45.1/CD45.2 recipient mice (n=4/condition) and contribution to platelets, erythrocytes, neutrophils **(f)** and bone marrow HSCs and progenitors **(g)** was monitored for 4 weeks (individual mice are shown). Transplanted LEK cells served as a positive control (mean and SEM, n=3). Transplanted B6.CD45.1/.2 carrier splenocytes served as negative controls (n=3, not shown). **h**, Probability of physiological thrombopoiesis proceeding via the direct CD48^−/lo^ MkP pathway (mean and 95% confidence bounds).

**Extended Data Figure 7.**
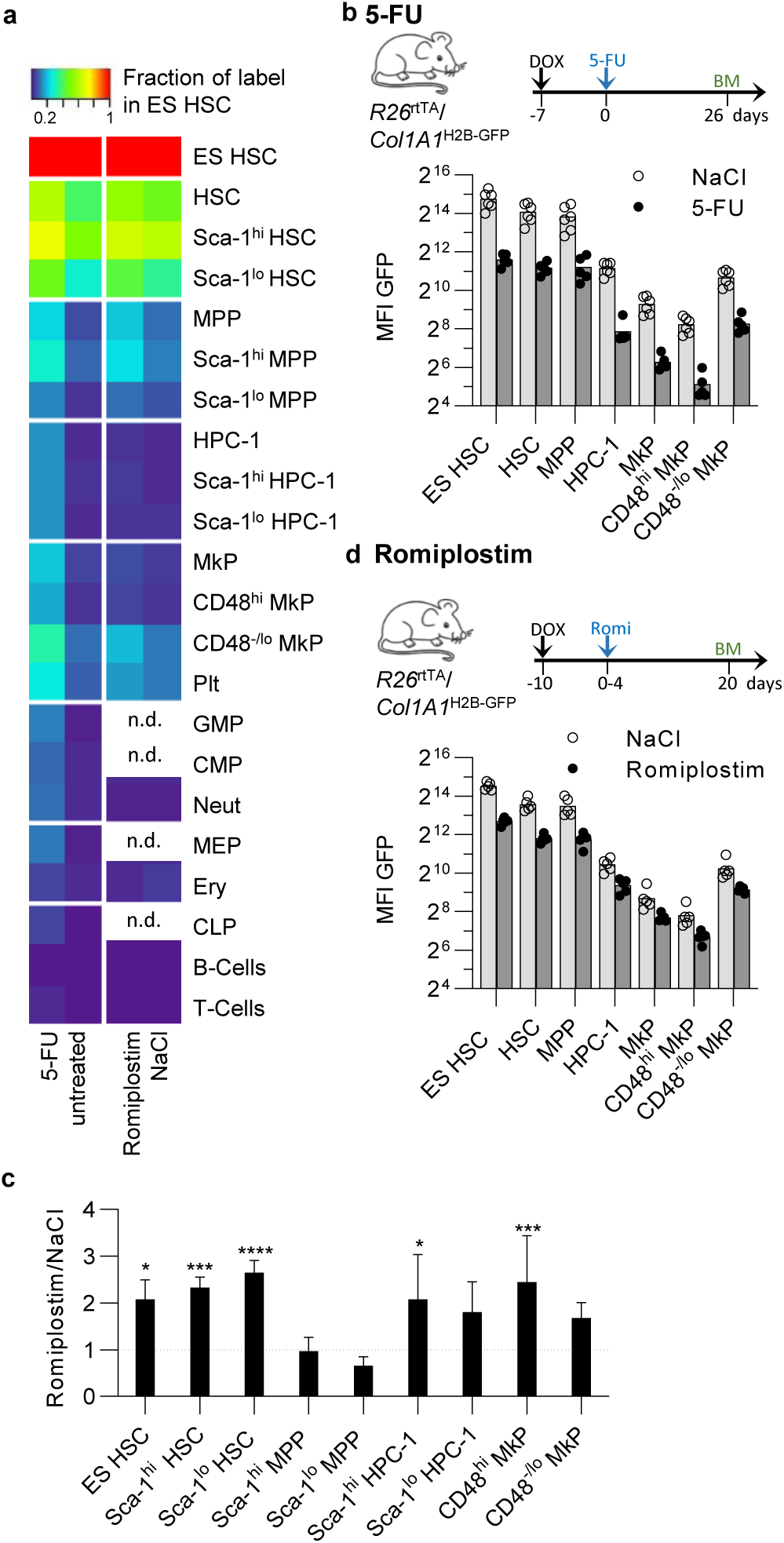
Perturbation of hematopoiesis by 5-FU and romiplostim. **a**, *Fgd5*^ZsGreen:CreERT2^*/R26*^LSL-tdRFP^ mice were TAM-induced and treated with 5-FU (left heat map, same animals as in Fig. 4a) or romiplostim (right heat map, same animals as in Fig. 4d). Heat map shows RFP labelling as fraction of label in ES HSC. **b**, *R26*^rtTA^/*Col1A1*^H2B-GFP^ animals were Dox-induced and treated either with 5-FU or saline (NaCl, n=6/condition). Cell populations were analysed for H2B-GFP retention at indicated time points. H2B-GFP mean fluorescence intensities (MFI) from individual mice and means thereof (bars) are shown in log_2_ display. Population-specific background H2B-GFP fluorescence was subtracted. **c**, Changes in bone marrow population size of *Fgd5*^ZsGreen:CreERT2^*/R26*^LSL-tdRFP^ mice after romiplostim treatment compared to saline (NaCl, dotted line) treated controls (mean + SEM, one-way ANOVA with Sidak correction, same animals as in Fig. 4d). Absolute numbers of HSPCs were normalized to body weight. **d**, *R26*^rtTA^/*Col1A1*^H2B-GFP^ animals were Dox-induced and treated either with romiplostim or saline (NaCl, n=5/condition). Normalisation and display of data as in **b**.

**Extended Data Figure 8.**
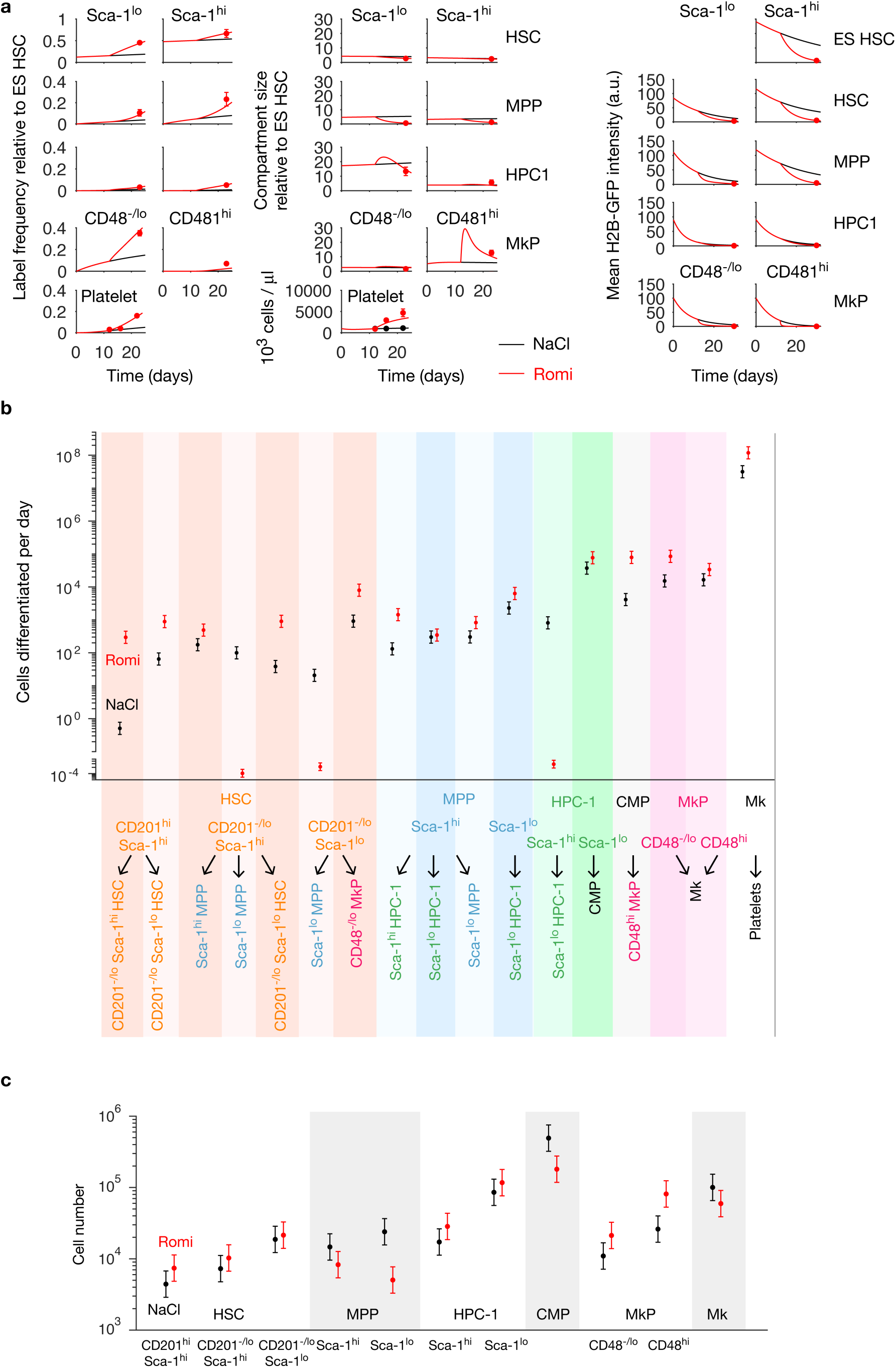
Modeling of romiplostim-enhanced thrombopoiesis. **a**, Model fitting of RFP label propagation (left), compartment size (middle) and H2B-GFP dilution (right) dynamics in unperturbed (NaCl, black) or romiplostim-treated (red) conditions. **b**, Inferred cell fluxes for all differentiation steps. **c**, Inferred cell counts of all compartments. In b and c, dots are for best-fit parameters and error bars indicate 95% prediction bands.

## Supplementary information

### Supplementary methods

#### 1 Mathematical modeling and inference of lineage pathways

##### 1.1 Dynamics of hematopoietic cell populations

The number of cells in a given hematopoietic population are determined by the balance of cell gain and loss. Cells are gained by influx through differentiation of progenitors and proliferation of cells in the given population; loss occurs by onward differentiation and death. Specifically, all stem and progenitor cells proliferate, whereas mature cells may not proliferate (e.g., erythrocytes or platelets). With the exception of tip stem cells (i.e., ES HSCs), all cell populations receive influx. All cell types, including HSCs ^1^, may die.

The mathematical balance equations for the cell numbers are based on the principle that the flux of cells—the number of cells undergoing a certain process (e.g., cell division) per unit time—is proportional to the number of cells that can potentially undergo this process. The proportionality factor is a rate constant that is the inverse of the average waiting time before a cell commits to the process in question (e.g., the average cell cycle time in the case of cell division). Hence, the balance equations are exact in principle. Their precision in describing the actual biology depends on the appropriate definition of cell populations. Here, we use a refined definition of the most upstream hematopoietic populations, HSCs, MPPs and HPC-1, that includes, in addition to the usual markers (LSK, CD48 and CD150) also CD201 (EPCR) for HSCs and Sca-1 surface level for HSCs, MPPs and HPC-1. Thus we go well beyond the resolution of previous studies ^2,3^.

Specifically, HSCs and progenitors can undergo self-renewal (with rate *σ*), divide asymmetrically (with rate *γ*), divide into two more differentiated daughter cells (with rate *ρ*, symmetric differentiating division), differentiate directly (with rate *μ*) or die (with rate *‹*) (Supplementary Fig. 1a). Based on these fundamental processes, the population dynamics for a linear lineage pathway (i.e., each product has exactly one precursor) are described by the following system of ordinary differential equations:

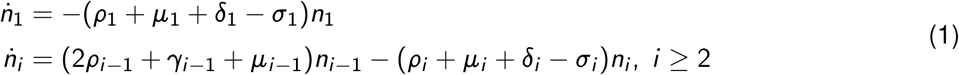

where population 1 are the tip stem cells (ES HSCs) and populations 2, 3,… are successive downstream populations. These equations can be rewritten in a simpler form:

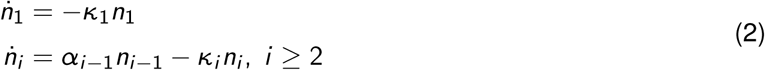

using two aggregate parameters:

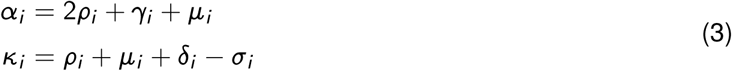

where *α*_*i*_ and *κ*_*i*_ are the total differentiation rate and net cell loss rate of compartment *i*, respectively. Based on this generic model, in the following sections we develop a quantitative framework of inferring differentiation pathways from the combination of HSC fate-mapping and cell proliferation data.

##### 1.2 Propagation of fate-mapping label

Let *m*_*i*_ and *n*_*i*_ be the labeled and total cell number of population *i*. The labeling frequency is defined by 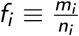. The dynamics of the labeling frequency is governed by,

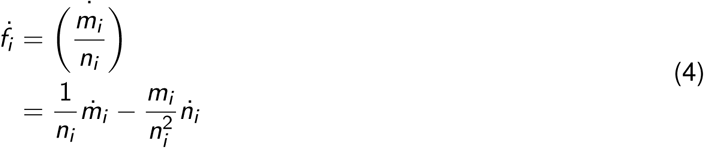

Assuming that heritable labeling (e.g., via RFP expression from the Rosa26 locus) does not alter cell proliferation, differentiation and death rates, we combine Equation 4 with Equation 2 and obtain

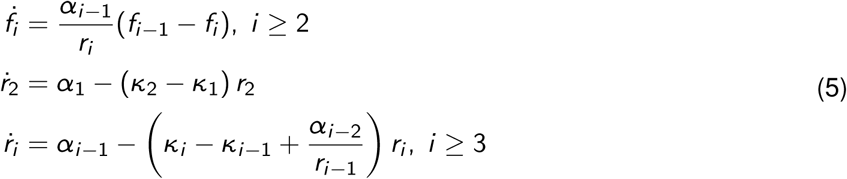

where for the tip stem cells *f*_1_ = constant, as shown experimentally (Figure 1d). The new variable *r*_*i*_ denotes the size ratio between neighboring compartments: 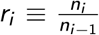^4^. The generic model can be readily extended to describe more complex lineage trees, e.g. with diverging and converging branches.

##### 1.3 Modeling H2B-GFP dynamics

To infer cell proliferation rates from the H2B-GFP dilution data, we model the temporal changes of the mean H2B-GFP intensity (total H2B-GFP intensity divided by cell number) in each cell population. The cell number dynamics is given by Equation 2, so next we derive the equations for total H2B-GFP intensity in each compartment.

We assume that H2B-GFP turnover and basal production change the fluorescence intensity of all the cell populations uniformly, whereas cell proliferation, differentiation and death do so in a population-specific manner. Let *X*_*i*_ (*t*) be the total H2B-GFP intensity of compartment *i* at time *t*. During time increment ∆*t*, H2B-GFP turnover and basal production cause the change in total intensity by,

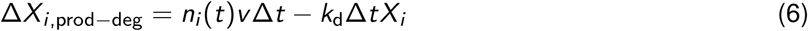

where *n*_*i*_ (*t*) is the cell number of compartment *i*, *v* the basal H2B-GFP production rate per cell and *k*_d_ the turnover rate.

Symmetric self-renewal redistributes existing H2B-GFP into more cells in the same population, thus altering the relative (i.e., per cell) but not the total H2B-GFP intensity of the population. By contrast, cell death or differentiation into the next population, i.e. *i* → *i* + 1, cause H2B-GFP loss from population *i*. The intensity loss equals the number of cells that leave the compartment times the H2B-GFP intensity taken away per cell. For example, the cell number loss after ∆*t* is *γ*_*i*_ *n*_*i*_ (*t*)∆*t* due to asymmetric differentiation. The H2B-GFP level loss per cell is half of the current level, i.e. 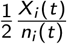 because only one of the two daughter cells goes to the next population. The change in total H2B-GFP level is then 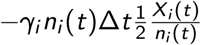. For cells undergoing death, symmetric or direct differentiation, all H2B-GFP of these cells is lost from population *i*. Thus, the total H2B-GFP loss after ∆*t* for compartment *i* is,

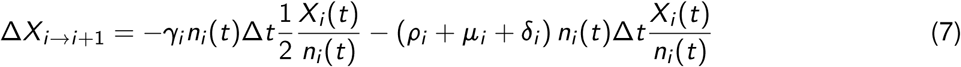

Furthermore, each population, except for the tip stem cells, receives H2B-GFP from its direct upstream population via differentiation. In a similar way, we formulate this effect as,

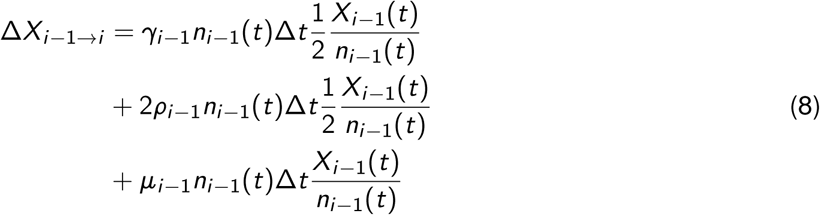

Therefore, the total H2B-GFP level of compartment *i* at time *t* + ∆*t* is,

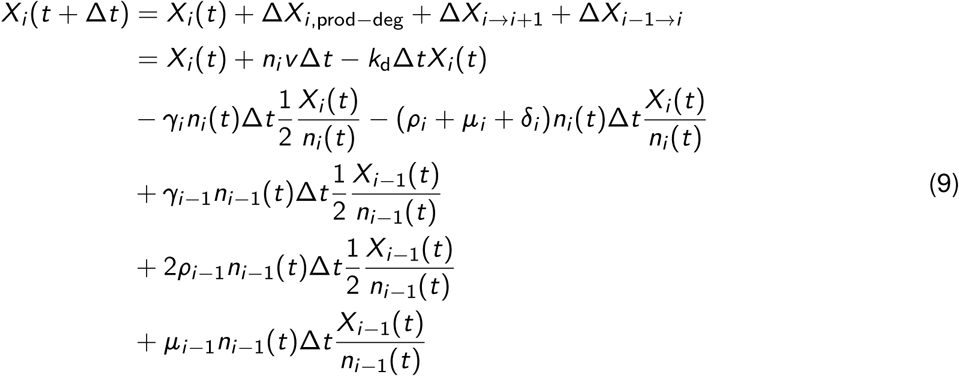

The dynamics of the mean H2B-GFP intensity 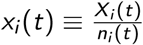 obeys,

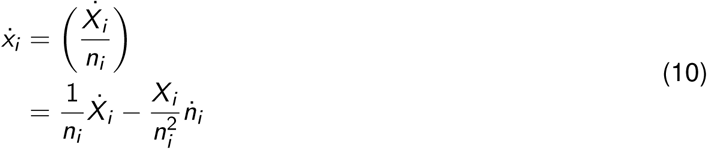

Combining this with Equation 1, we obtain

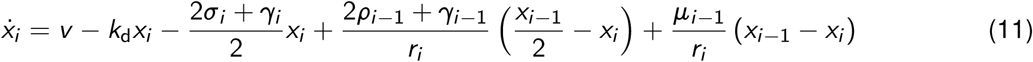

This equation shows how each elementary process contributes to the H2B-GFP dynamics. First, cell death, symmetric and direct differentiation of a given population *i* do not affect its own mean H2B-GFP level (none of the parameters *δ*_*i*_, *ρ*_*i*_ or *μ*_*i*_ appear in the equation). Second, H2B-GFP intensity is diluted by symmetric self-renewal and asymmetric differentiation. Third, cells entering via differentiation from the upstream population can either increase or reduce the mean H2B-GFP level, depending on the intensity difference. Finally, H2B-GFP dilution is also determined by its own degradation (with *k*_d_). For all parameter inferences in this paper, we fixed *k*_d_ to its previously determined value in murine HSCs, corresponding to a half-life of 42 days ^5^, thus improving inference of the relevant rate constants of HSC and progenitor proliferation, differentiation and death.

We rewrite Equation 11 using the aggregate parameters defined in Equation 3

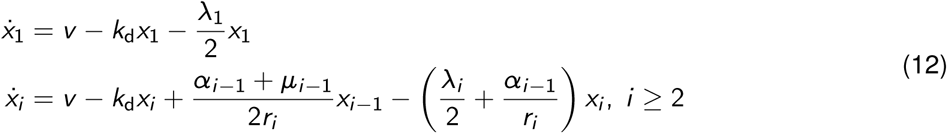

and introduce another aggregate parameter λ_*i*_, the total self-renewal rate

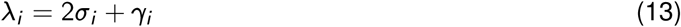

In summary, we obtain a system of differential equations for cell numbers (Equation 2), frequencies of fate-mapping label (Equation 5) and average H2B-GFP intensities (Equation 12) in hematopoietic cell populations. The key parameters characterizing the dynamics of cell proliferation and differentiation—total differentiation rates *α*_*i*_, direct differentiation rates *μ*_*i*_, net loss rates *κ*_*i*_ and total self-renewal rates *λ*_*i*_ μcan all be time-dependent (e.g., change with increasing age of the animal).

##### 1.4 Demonstrating inference of lineage pathways

Label propagation and H2B-GFP dilution experiments yield, respectively, differentiation and proliferation rates of stem and progenitor cells. We therefore reasoned that the combination of the two types of experimental data will allow inference of both differentiation and proliferation rates in a complex lineage tree. We have advanced this argument recently for the combination of HSC fate mapping and BrdU incorporation (instead of H2B-GFP dilution) ^4^. The principal new aspect in this paper is that we show that combining these types of experimental data also yields information on the topology of the underlying lineage tree. To show this, we first developed the idea with a toy model (Extended Data Figure 3). To this end, we simulated data for a known lineage topology, and then asked whether model selection based on these *in silico* data with a broader set of possible lineage topologies indeed identifies the correct topology (ground truth).

We collect the equations from Equation 5 and 12,

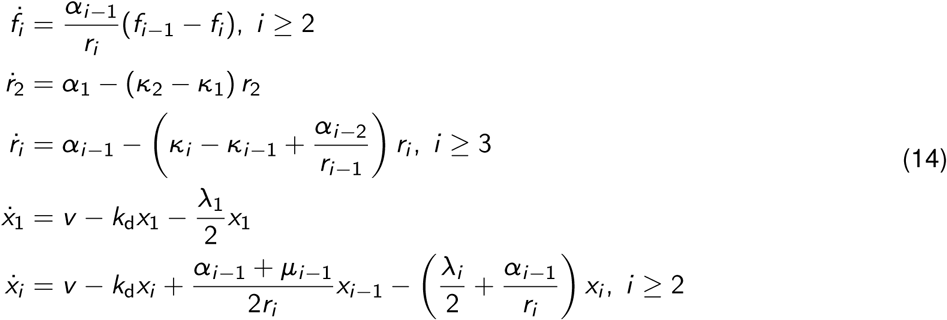

where

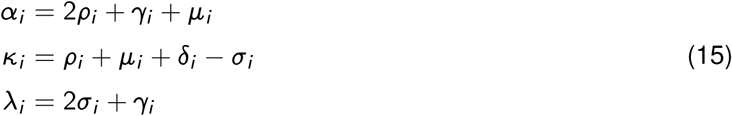

The value of total net cell loss *κ*_*i*_ can be positive or negative. To be more practical, we substitute *κ*_*i*_ with other naturally positive parameters,

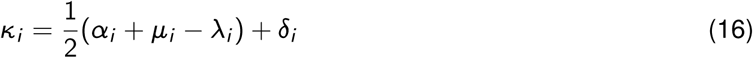

According to Equation 14, each compartment is governed by 4 parameters (*α*_*i*_, *λ*_*i*_, *μ*_*i*_ and *δ*_*i*_), while there are 5 underlying elementary parameters (*ρ*_*i*_, *γ*_*i*_, *σ*_*i*_, *μ*_*i*_ and *δ*_*i*_). Thus, model inference has a natural limitation in identifying each individual fundamental parameters. However, inference based on the aggregate parameters is sufficient for identifying differentiation pathways.

Next, we test this framework by inferring differentiation pathways based on in silico data. We consider a lineage tree with one stem cell (S), two progenitor cells (P1 and P2) and three mature cells (M1, M2 and M3) with a unique pathway topology (Extended Data Fig. 3b) and assign physiologically reasonable parameters. We obtained the in silico data by adding white noise to the simulated data (label propagation, compartment size and H2B-GFP dilution data), with different coefficients of variance for different time points. We enumerated 6 possible model schemes (without lineage convergence), and performed statistical model selection based on the simulated data (Extended Data Fig. 3c). The equations for the true model scheme are:

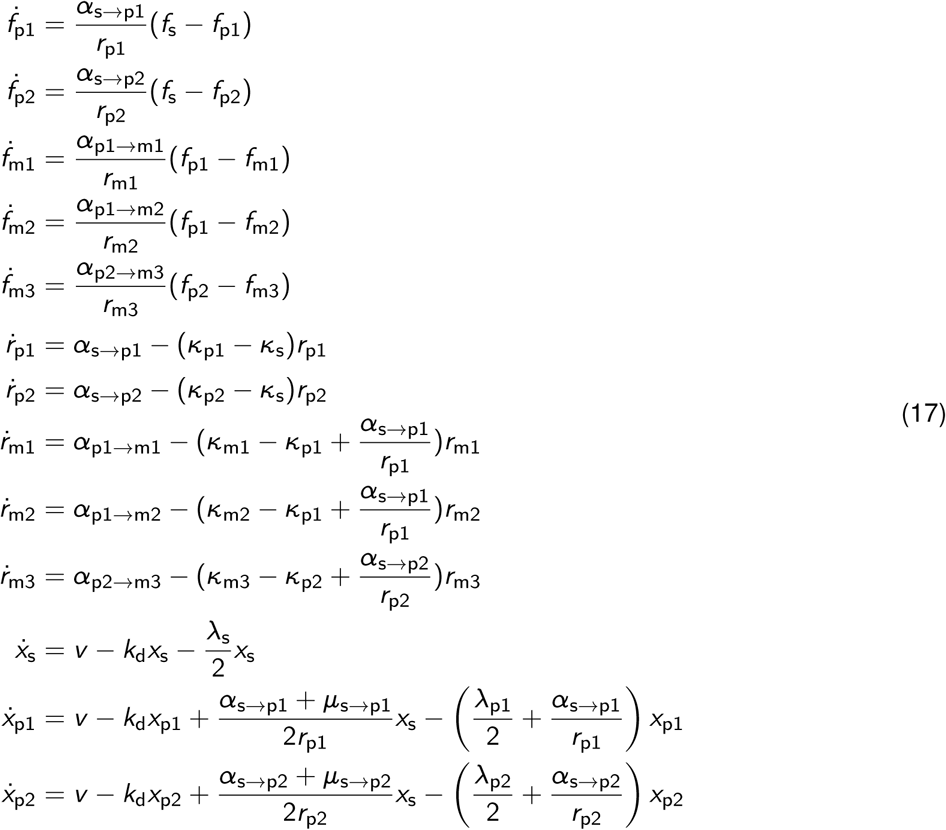

where we substitute the net cell loss rates *κ* of the stem and progenitor cells using the following equations,

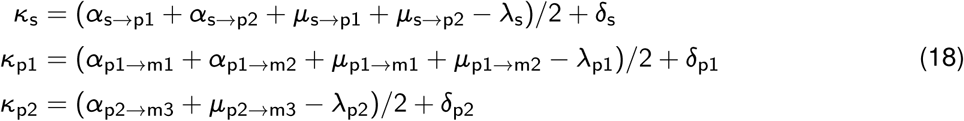

Here, we do not explicitly consider the proliferation or death of the mature cells but simply quantify their net loss rate (in the case of non-proliferating mature cells, the death rate will equal the net loss rate).

The model selection successfully discovered the true model (Extended Data Fig. 3d and e). In stark contrast, the true model could not be singled out by inference without the proliferation data (Extended Data Fig. 3f). To conclude, combining fate-mapping and cell proliferation data allows both identification of lineage topology and quantification the cell fluxes in the lineage pathways.

#### 2 Inferring lineage pathways of native hematopoiesis from experimental data

##### 2.1 Modeling lineage pathways of hematopoietic stem and progenitor cells

To address the experimental data on HSC fate mapping, augmented by tracking of mitotic history of HSCs and progenitors via H2B-GFP, we devised a family of models, with each HSC and progenitor population divided into Sca-1 high and Sca-1 low subpopulations (Supplementary Fig. 1b). HSC subpopulations are further divided by CD201 level. The CD201^−/lo^ Sca-1^lo^ subpopulation has very few cells, so it is not considered in the models. To identify the differentiation pathways, we performed statistical model selection against the experimental data. The equations for the full model with all possible differentiation and Sca-1 transition steps read:

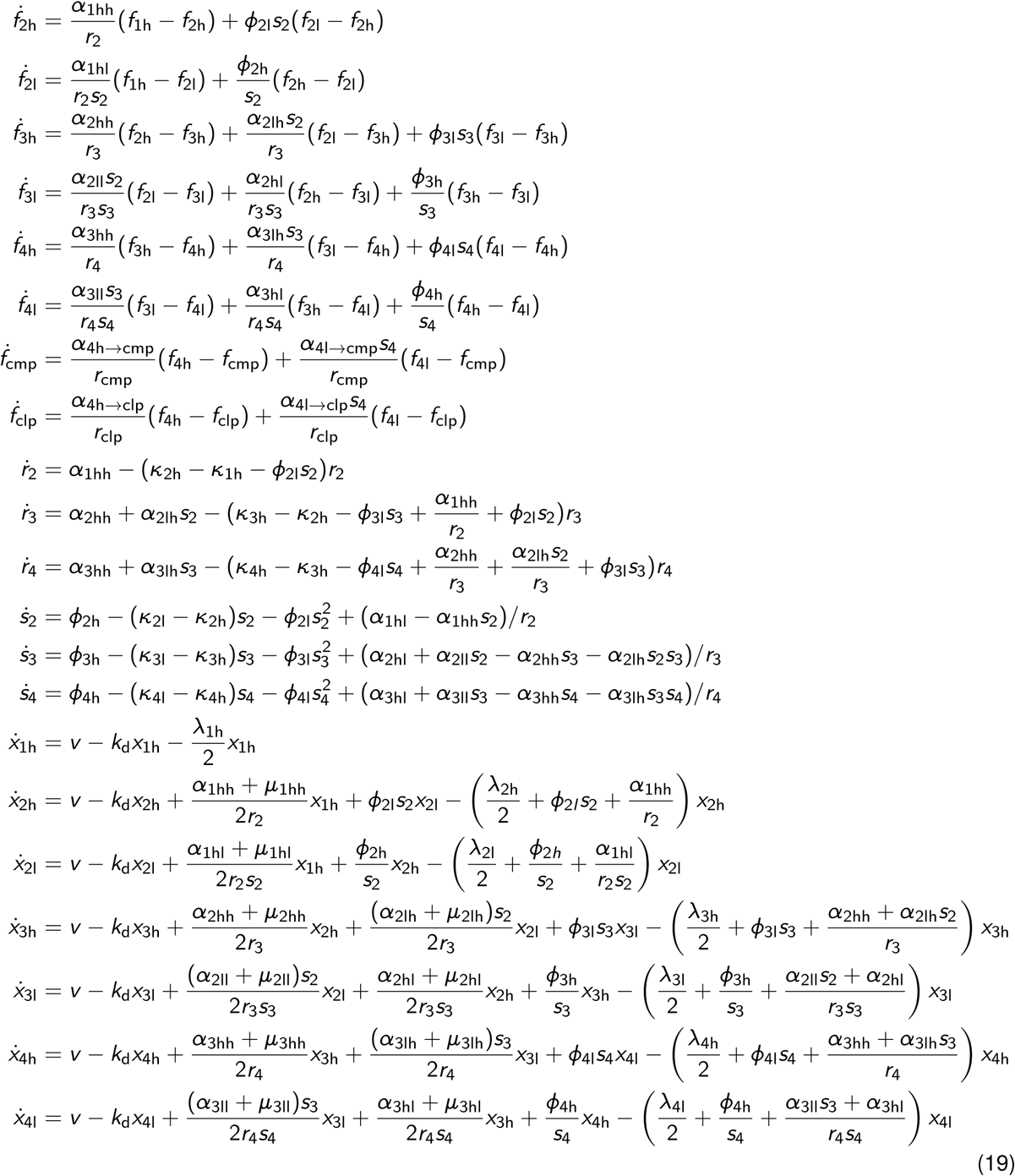

where

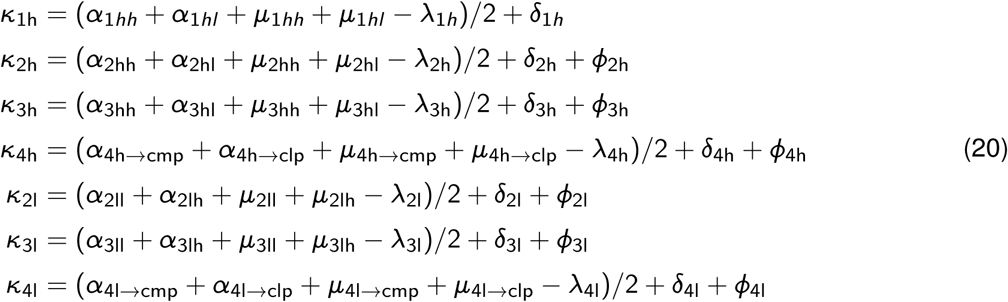

The subscript *i* in the model variables and parameters ranges from 1 to 4, representing cell compartments CD201^hi^ HSC, CD201^−/lo^ HSC, MPP and HPC-1, respectively. In addition, the subscripts *i*h and *i*l stand for compartment *i* Sca-1 high and Sca-1 low, respectively. The variable *r*_*i*_ is the compartment size ratio between two neighboring Scal-1^hi^ compartments, i.e. *r*_*i*_ = *n*_*i*h_/*n*_*i*−1h_; *i* = 2, 3, 4. *s*_*i*_ is the size ratio between Sca-1^lo^ and Sca-1^hi^ sub-compartments at each level, i.e. *s*_*i*_ = *n*_*i*l_/*n*_*i*h_, *i* = 2, 3, 4. The parameter *φ* is the transition rate between Sca-1^hi^ and Sca-1^lo^ subpopulations. Detailed descriptions of all parameters are given in Supplementary Table 1.

Model selection is done with a comprehensive array of sub-models of the most general model, thus explicitly introducing parsimony and allowing model ranking via the Akaike information criterion. The equations for any sub-model are derived from Equation 19 by setting specific parameters to zero. Initial fitting results show the most direct differentiation rates *μ* tend to be zero, implying that asymmetric cell division (possibly augmented by symmetric differentiating division) is the primary mode of differentiation. Hence we set all *μ*_*i*_ to zero for simplicity. We fit all the models to the *Fgd5*^*ZsGreen:CreERT2*^ label propagation and H2B-GFP dilution data and ranked the models using the bias-corrected Akaike information criterion (AICc). The top ranked models show two common features: first, parallel differentiation via Sca-1^hi^ and Sca-1^lo^ paths; second, irreversible transition from Sca-1^hi^ to Sca-1^lo^ states. Models bearing one or neither of the two features ranked poorly.

##### 2.2 Quantifying the probability of lineage pathways

There is one single pathway for lymphoid lineage, where Sca-1 level is consistently high from tip HSC to HPC-1. On the other hand, myeloid cells can be derived via 6 pathways. To quantify the contributions of these pathways, we calculated pathway probability based on differentiation history. We started at the final cell state *z*_*i*_, and calculated the probability of its ancestral line of cells to have differentiated via a pathway *z*_0_ → *z*_1_ → … → *z*_*i*_. To do this, we back-tracked from state *z*_*i*_ and denoted the probability to have arrived from the precursor state *z*_*i*−1_ by *p*(*z*_*i*−1_ → |*z*_*i*_). Because we consider a Markov state network, the differentiation decision at each state is independent of the differentiation history, such that the pathway probability factorizes:

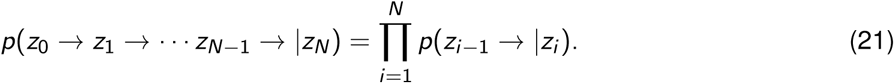

When an expanding cell population reaches a state of steady growth, the elementary transition probability can be written as:

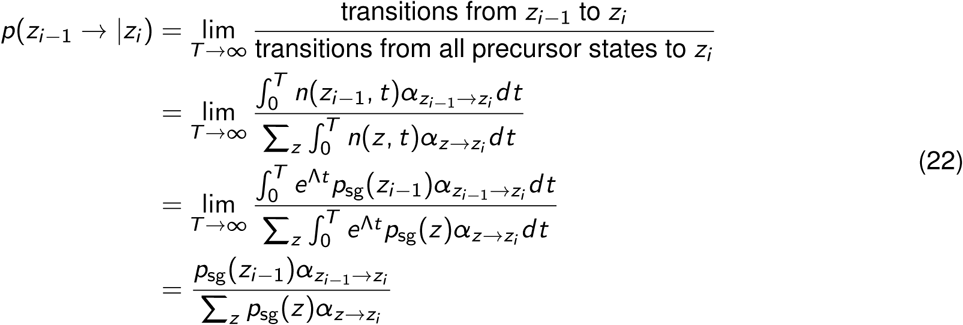

where the sums run over all possible precursor states; Λ is the dominant eigenvalue of the dynamic system, that is, the population growth rate; *p*_sg_ (*z*_*i*_) is the relative compartment size of *z*_*i*_ to the sum of all populations at steady growth; and *α* is the differentiation rate. Combining Equations 21 and 22, we calculated the probability of all six paths from tip HSC to CMP at steady growth.

##### 2.3 Model inference including mature cell populations

We extend the current model to describe differentiation pathways of mature blood cells, with a focus on platelets. It is known that platelets arise from conventional CMP-MkP pathway. However, the labeling frequency of platelets is higher than that of MPP, which suggests that the conventional pathway is not the only route to produce platelets. We therefore extended the model to allow MkP to directly receive cells from CD201^−/lo^ Sca-1^lo^ HSCs, and fit the model to the collective data of HSPC, MkP and platelets. However, the model fails to capture the label propagation data of MkP and platelets, because the measured MkP labeling frequency is still lower than platelets (Figure 3c). A potential solution to this controversy is to assume two, in the simplest case, heterogeneous subpopulations in MkP, one of which is produced directly from HSC (Extended Data Fig. 6a). The model of heterogeneous MkP compartment is supported by the data (Extended Data Fig. 6b).

CD48^−/lo^ MkP arises directly from Sca-1^lo^ HSC, while CD48^hi^ MkP is produced via the conventional CMP pathway. Moreover, the two-compartment model faithfully captured the data of MkP subpopulations.

The corresponding model equations are:

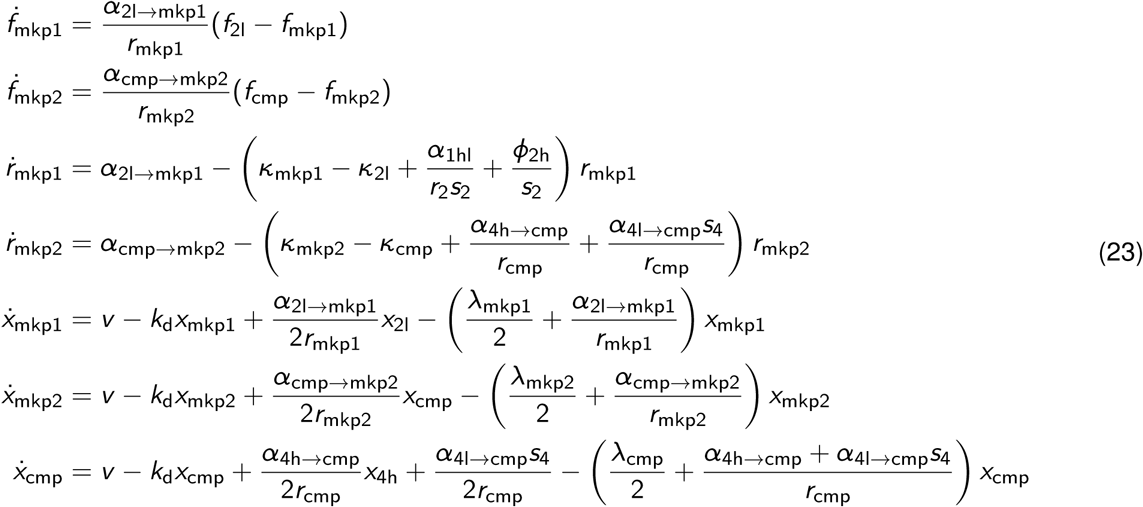

where mkp1 and mkp2 in the subscripts represent CD48^−/lo^ and CD48^hi^ MkP, respectively. The net cell loss of Sca-1^lo^ HSC (compartment 2l) also contains the direct differentiation path to MkP CD48^−/lo^,

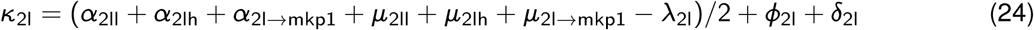

The equations for megakaryocytes and platelets are,

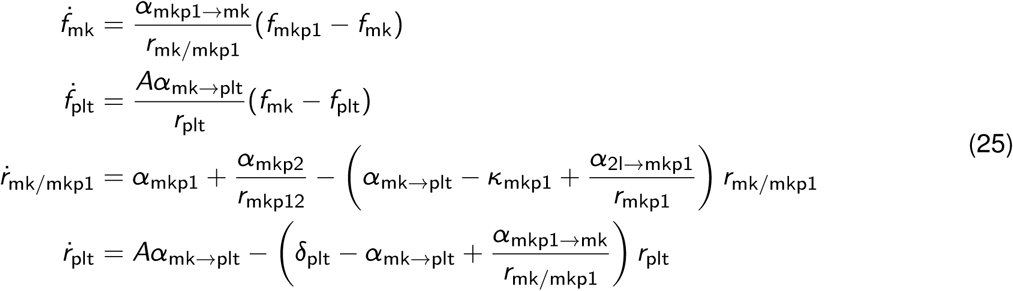

where

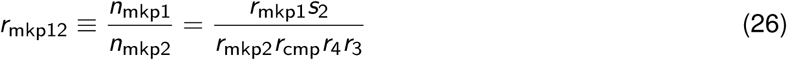

The model assumes no proliferation for megakaryocyte, and the cytoplasm of a megakaryocyte is shed into fragments, forming *A* = 10000 platelets ^6^, at rate *α*_mk→plt_.

We then use the best fit parameters to calculate the contribution of the two pathways to platelet production, which is determined by the fluxes (cell differentiated per day) via CD48^−/lo^ MkP and CD48^hi^ MkP.

##### 2.4 Rate changes upon romiplostim treatment

To quantify the effect of romiplostim on thrombopoiesis, we allowed the model parameters to change in a stepwise manner corresponding to the treatment timing. Up to 2 weeks, all parameters remain at their values for unperturbed native hematopoiesis. At this time point, we simulated romiplostim treatment by allowing a step-change of all parameters. Interestingly, the parameters along the CD48 low MkP pathway change most compared with the unperturbed condition (Supplementary Table 1), and romiplostim preferentially regulates thrombopoiesis via CD48^−/lo^ MkP pathway.

---

**Supplementary Fig. 1.**
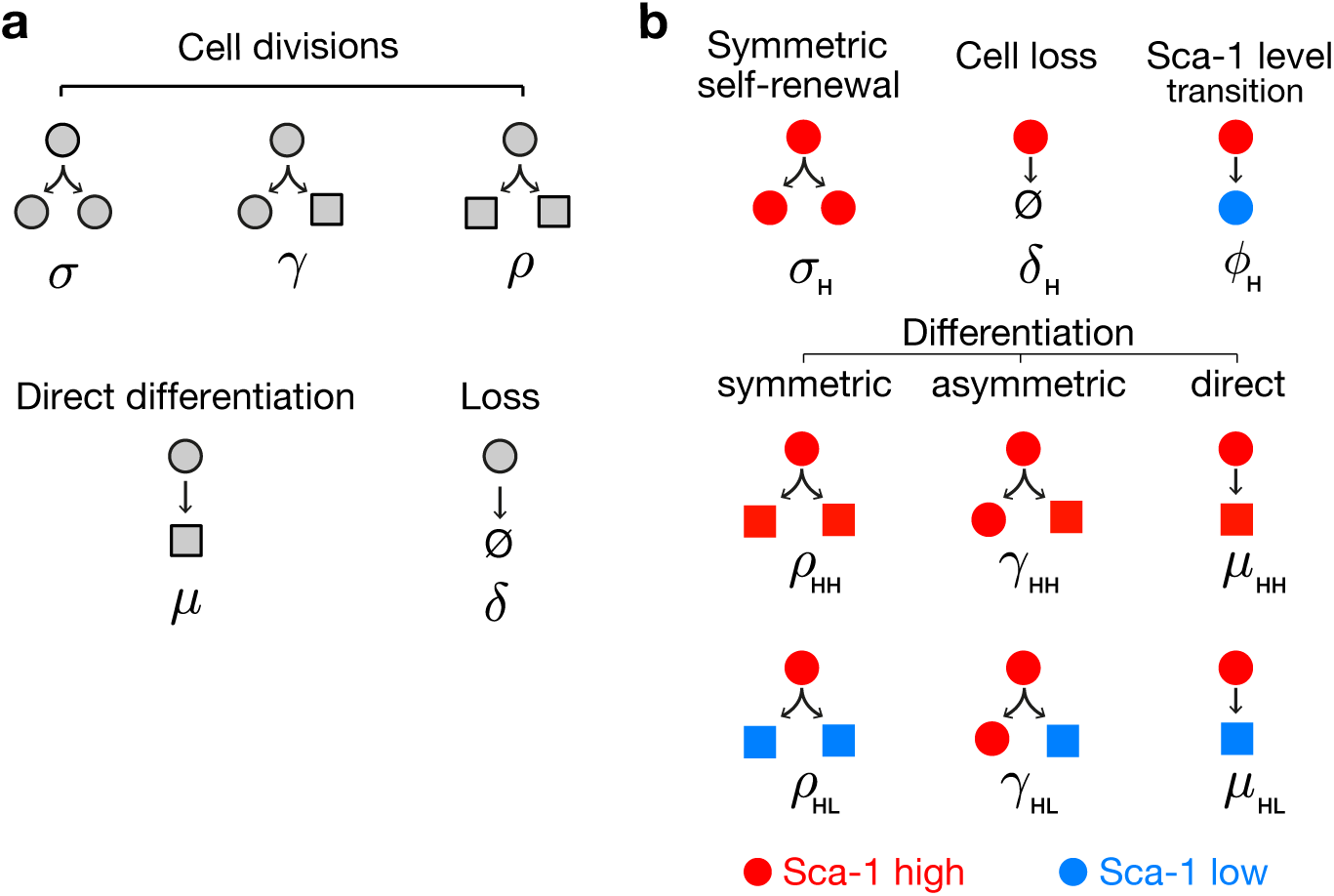
Elementary processes governing the numbers of stem and progenitor cells. (a) Fundamental cell fates of a stem or progenitor cell. (b) Extended scheme considering Sca-1^hi^ (red) and Sca-1^lo^ (blue) subpopulations. The elementary processes of a Sca-1^hi^ stem/progenitor cell are shown, which are symmetric with those of a Sca-1^lo^ cell. The subscripts H and L of the parameters stand for Sca-1^hi^ and Sca-1^lo^, respectively. HH and HL represent the differentiation without and with decline in Sca-1 level.

**Supplementary Table. 1.** Inferred parameter values for native hematopoiesis and activated hematopoiesis after romiplostim treatment.

| Name | Description | Best fit [95% confidence bounds] in units day<sup>-1</sup>,<br>unless specified otherwise |  |
| --- | --- | --- | --- |
|  |  | Unperturbed | TPO |
| $\alpha_{1hh}$ | Differentiation rate of<br>CD201<sup>hi</sup>Sca-1<sup>hi</sup> HSC →<br>CD201<sup>-/lo</sup>Sca-1<sup>hi</sup> HSC | 1e-4 [2.5e-5 2e-4] | 0.04 (0 0.1) |
| $\alpha_{1hl}$ | Differentiation rate of<br>CD201<sup>hi</sup>Sca-1<sup>hi</sup> HSC →<br>CD201<sup>-/lo</sup>Sca-1<sup>lo</sup> HSC | 0.013 [0.01 0.015] | 0.12 [0.07 0.15] |
| $\alpha_{2hh}$ | Differentiation rate of<br>CD201<sup>-/lo</sup>Sca-1<sup>hi</sup> HSC →<br>Sca-1<sup>hi</sup> MPP | 0.03 [0.02 0.04] | 0.05 [0.018 0.11] |
| $\alpha_{2hl}$ | Differentiation rate of<br>CD201<sup>-/lo</sup>Sca-1<sup>hi</sup> HSC →<br>Sca-1<sup>lo</sup> MPP | 0.013 [0.008 0.018] | 1e-8 (0 0.04) |
| $\phi_{2h}$ | Transition rate of<br>CD201<sup>-/lo</sup>Sca-1<sup>hi</sup> HSC →<br>CD201<sup>-/lo</sup>Sca-1<sup>lo</sup> HSC | 6e-4 (0 0.01) | 0.08 [0.05 0.17] |
| $\alpha_{2ll}$ | Differentiation rate of<br>CD201<sup>-/lo</sup>Sca-1<sup>lo</sup> HSC →<br>Sca-1<sup>lo</sup> MPP | 9e-4 [6e-4 0.001] | 1e-8 (0 0.019) |
| $\alpha_{2l \rightarrow mklp1}$ | Differentiation rate of<br>CD201<sup>-/lo</sup>Sca-1<sup>lo</sup> HSC →<br>CD48<sup>-/lo</sup> MkP | 0.05 [0.03 0.1] | 0.37 [0.29 0.47] |
| $\alpha_{3hh}$ | Differentiation rate of Sca-1<sup>hi</sup><br>MPP → Sca-1<sup>hi</sup> HPC-1 | 0.008 [0.006 0.01] | 0.16 [0.09 0.32] |
| $\alpha_{3hl}$ | Differentiation rate of Sca-1<sup>hi</sup><br>MPP → Sca-1<sup>lo</sup> HPC-1 | 3e-6 (0 0.006) | 0.05 (0 0.28) |
| $\phi_{3h}$ | Transition rate of Sca-1<sup>hi</sup> MPP<br>→ Sca-1<sup>lo</sup> MPP | 0.016 [0.013 0.017] | 0.09 (0 0.15) |
| $\alpha_{3ll}$ | Differentiation rate of Sca-1<sup>lo</sup><br>MPP → Sca-1<sup>lo</sup> HPC-1 | 0.15 [0.12,0.17] | 1.3 [0.5 2.1] |
| $\alpha_{4h \rightarrow clp}$ | Differentiation rate of Sca-1<sup>hi</sup><br>HPC-1 → CLP | 0.1 [0.08 0.15] | 0.4 [0.14 0.56] |
| $\phi_{4h}$ | Transition rate of Sca-1<sup>hi</sup><br>HPC-1 → Sca-1<sup>lo</sup> HPC-1 | 0.038 [0.014 0.059] | 1e-8 (0 0.13) |

**Supplementary Table. 1.** Inferred parameter values for native hematopoiesis and activated hematopoiesis after romiplostim treatment.

| Name | Description | Best fit [95% confidence bounds] in units day<sup>-1</sup>,<br>unless specified otherwise |  |
| --- | --- | --- | --- |
|  |  | Unperturbed | TPO |
| $\alpha_{4l \rightarrow cmp}$ | Differentiation rate of Sca-1<sup>lo</sup><br>HPC-1 → CMP | 0.4 [0.35 0.45] | 0.64 [0.5 0.9] |
| $\alpha_{cmp \rightarrow mkp2}$ | Differentiation rate of CMP →<br>CD48<sup>hi</sup> MkP | 0.008 [0.005 0.015] | 0.4 [0.3 0.6] |
| $\alpha_{mkp1 \rightarrow mk}$ | Differentiation rate of CD48<sup>-/lo</sup><br>MkP → Mk | 1.4 [0.9,1.9] | 4 [3.45 4) |
| $\alpha_{mkp2 \rightarrow mk}$ | Differentiation rate of CD48<sup>hi</sup><br>MkP → Mk | 0.64 [0.41 0.85] | 0.40 [0.26 0.6] |
| $\alpha_{mk \rightarrow plt}$ | Differentiation rate of Mk →<br>platelets | 0.3 [0.08 0.57] | 1.95 [1.5 2.5] |
| $\lambda_{1h}$ | Self-renewal rate of<br>CD201<sup>hi</sup>Sca-1<sup>hi</sup> HSC | 0.022 [0.020 0.023] | 0.27 [0.26 0.28] |
| $\lambda_{2h}$ | Self-renewal rate of<br>CD201<sup>-/lo</sup>Sca-1<sup>hi</sup> HSC | 0.079 [0.076 0.083] | 0.27 [0.13 2.2] |
| $\lambda_{2l}$ | Self-renewal rate of<br>CD201<sup>-/lo</sup>Sca-1<sup>lo</sup> HSC | 0.096 [0.088 0.11] | 0.31 [0.20 0.46] |
| $\lambda_{3h}$ | Self-renewal rate of Sca-1<sup>hi</sup><br>MPP | 0.03 [0.026 0.033] | 0.20 [0.02 0.6] |
| $\lambda_{3l}$ | Self-renewal rate of Sca-1<sup>lo</sup><br>MPP | 0.13 [0.12 0.15] | 0.78 [0.19 1.45] |
| $\lambda_{4h}$ | Self-renewal rate of Sca-1<sup>hi</sup><br>HPC-1 | 0.18 [0.16 0.20] | 0.33 [0.09 3.3] |
| $\lambda_{4l}$ | Self-renewal rate of Sca-1<sup>lo</sup><br>HPC-1 | 0.32 [0.28 0.36] | 0.47 [0.32 0.82] |
| $\lambda_{cmp}$ | Self-renewal rate of CMP | 0.35 [0.05,3.7] | 3.7 [0.23 4) |
| $\lambda_{mkp1}$ | Self-renewal rate of CD48<sup>-/lo</sup><br>MkP | 0.13 [0.02 0.56] | 1.26 [0.89 1.74] |
| $\lambda_{mkp2}$ | Self-renewal rate of CD48<sup>hi</sup><br>MkP | 1.5e-7 (0 0.29] | 2e-8 (0 0.18] |
| $\kappa_{clp}$ | Net cell loss rate of CLP | 0.016 [0.01 0.02] | fixed value as<br>unperturbed |

**Supplementary Table. 1.** Inferred parameter values for native hematopoiesis and activated hematopoiesis after romiplostim treatment.

| Name | Description | Best fit [95% confidence bounds] in units day<sup>-1</sup>,<br>unless specified otherwise |  |
| --- | --- | --- | --- |
|  |  | Unperturbed | TPO |
| $\delta_{1h}$ | Death rate of CD201<sup>hi</sup>Sca-1<sup>hi</sup> HSC | 1e-08 (0 1e-3] | fixed value as unperturbed |
| $\delta_{2h}$ | Death rate of CD201<sup>-/lo</sup>Sca-1<sup>hi</sup> HSC | 0.03 [0.017 0.035] | fixed value as unperturbed |
| $\delta_{2l}$ | Death rate of CD201<sup>-/lo</sup>Sca-1<sup>lo</sup> HSC | 0.023 (0,0.04] | fixed value as unperturbed |
| $\delta_{3h}$ | Death rate of Sca-1<sup>hi</sup> MPP | 1e-8 (0 0.001] | fixed value as unperturbed |
| $\delta_{3l}$ | Death rate of Sca-1<sup>lo</sup> MPP | 1e-8 (0 0.01] | fixed value as unperturbed |
| $\delta_{4h}$ | Death rate of Sca-1<sup>hi</sup> HPC-1 | 3e-8 (0 0.01] | fixed value as unperturbed |
| $\delta_{4l}$ | Death rate of Sca-1<sup>lo</sup> HPC-1 | 2e-8 (0 0.02] | fixed value as unperturbed |
| $\delta_{cmp}$ | Death rate of CMP | 1e-06 (0 0.97] | fixed value as unperturbed |
| $\delta_{mkp1},$<br>$\delta_{mkp2}$ | Death rate of MkP | 1e-6 (0 0.2] | fixed value as unperturbed |
| $\nu$ | Basal production of H2B-GFP | 0.0016 [0.0015 0.0018]<br>(a.u. per day) | fixed value as unperturbed |
| $k_d$ | H2B-GFP turnover rate | $\ln(2)/42 = 0.0165$ | fixed value<sup>5</sup> |

**Supplementary Table 2.** Antibodies for flowcytometry.

Bone marrow HSPCs stainings
| Antibody | Fluorochrome | Clone | Isotope | Supplier |
| --- | --- | --- | --- | --- |
| CD34 | eF660 | RAM34 | Rat IgG2a, κ | eBioscience |
| CD34 | eF450 | RAM34 | Rat IgG2a, κ | eBioscience |
| CD41 | BV605 | MWReg30 | Rat IgG1, κ | Biolegend |
| CD45.1 | APC | A20 | Mouse IgG2a, κ | Biolegend |
| CD45.2 | FITC | 104 | Mouse IgG2a, κ | Biolegend |
| CD45.2 | Brilliant Violet 711 | 104 | Mouse IgG2a, κ | Biolegend |
| CD48 | Brilliant Violet 421 | HM48-1 | Ar. Hamster IgG | BD |
| CD48 | PE | HM48-1 | Ar. Hamster IgG | eBioscience |
| CD117 (c-Kit) | APC/eF780 | 2B8 | Rat IgG2b, κ | eBioscience |
| CD127 | PE/Cy7 | A7R34 | Rat IgG2a, κ | Biolegend |
| CD135 | BV421 | A2F10 | Rat IgG2a, κ | Biolegend |
| CD150 | PE/Cy7 | TC15-12F12.2 | Rat IgG2a, λ | Biolegend |
| CD16/32 | Alexa Fluor 700 | 93 | Rat IgG2a, λ | eBioscience |
| CD201 (EPCR) | APC | eBio1560 | Rat IgG2b, κ | eBioscience |
| Ly-6A/E (Sca-1) | PerCP/Cy5.5 | D7 | Rat IgG2a, κ | eBioscience |

Peripheral blood stainings
| Antibody | Fluorochrome | Clone | Isotope | Supplier |
| --- | --- | --- | --- | --- |
| CD3e | APC | 145-2C11 | Ar. Hs. IgG | eBioscience |
| CD3e | PerCP/eF710 | eBio500A2 | Syr.Hs. IgG | eBioscience |
| CD3e | APC | 145-2C11 | Ar. Hs. IgG | eBioscience |
| CD11b | APC/eF780 | M1/70 | Rat IgG2b, κ | eBioscience |
| CD11b | FITC | M1/70 | Rat IgG2b, κ | eBioscience |
| CD19 | PE/Cy7 | eBio1D3 | Rat IgG2a, κ | eBioscience |
| CD41 | APC | eBioMWReg30 | Rat / IgG1, κ | eBioscience |
| CD45.1 | APC | A20 | Mouse IgG2a, κ | eBioscience |
| CD45.2 | FITC | 104 | Mouse IgG2a, κ | Biolegend |
| Ly-6C/G (Gr1) | eF450 | RB6-8C5 | Rat IgG2b, κ | eBioscience |
| Ly-6C/G (Gr-1) | PerCP/Cy5.5 | RB6-8C5 | Rat IgG2b, κ | eBioscience |
| Ter-119 | FITC | TER-119 | Rat IgG2b, κ | Biolegend |

Biotinylated Antibodys
| Antibody | Fluorochrome | Clone | Isotope | Supplier |
| --- | --- | --- | --- | --- |
| CD3e | Biotin | 145-2C11 | Ar. Hs. IgG | eBioscience |
| CD4 | Biotin | GK1.5 | Rat IgG2b, κ | eBioscience |
| CD8a | Biotin | 53-6.7 | Rat IgG2a, κ | eBioscience |
| CD11b | Biotin | M1/70 | Rat IgG2b, κ | eBioscience |
| CD19 | Biotin | eBio1D3 | Rat IgG2a, κ | eBioscience |
| CD45R (B220) | Biotin | RA3-6B2 | Rat IgG2a, κ | Biolegend |
| Ly-6C/G (Gr-1) | Biotin | RB6-8C5 | Rat IgG2b, κ | eBioscience |
| NK-1.1 | Biotin | PK136 | IgG2a, κ | eBioscience |
| Ter-119 | Biotin | TER-119 | Rat IgG2b, κ | Biolegend |

Secondary Reagents
| Secondary Reagent | Fluorochrome | Supplier |
| --- | --- | --- |
| Streptavidin | V500 | BD |
| Streptavidin | eFluor710 | eBioscience |

